# Alternative EEG pre-processing pipelines can lead to conflicting conclusions regarding cortical excitation/inhibition ratio

**DOI:** 10.1101/2024.11.08.622698

**Authors:** Frigyes Samuel Racz, Diego Mac-Auliffe, Peter Mukli, John Milton, Juan Luis Cabrera, José del R. Millán

## Abstract

Confluent recent evidence indicates that the spectral slope of 1*/f* neurophysiological recordings is correspondent to cortical excitation/inhibition (E/I) ratio. In this framework, a steeper power spectrum (i.e., one with a larger spectral exponent *β*) is indicative of stronger inhibitory tone and thus lower E/I ratio, and vice versa. While the tools commonly utilized for estimating *β* are mostly consistent, there appears to be a lack of standardization among data processing protocols for slope analysis. In this work our goal is to draw attention to a fundamental consequence of this issue, namely that even in a confined, comparative research environment, applying different pre-processing steps to electroencephalography (EEG) data can lead to conflicting conclusions in terms of the E/I ratio. To this end, we analyzed resting-state EEG recordings in two independent datasets, containing data collected with eyes open (EO) and eyes closed (EC), with the latter considered as a physiological state with stronger inhibitory tone. Our analyses confirmed consistently in both cohorts that applying different spatial filtering schemes in an otherwise identical analytical pipeline indicated a decrease in E/I ratio over the prefrontal cortex in one case, but not the other when transitioning from EO to EC. In contrast, this same pattern was apparent over the occipital cortex regardless of the pre-processing scheme. This empirical evidence calls for the development of a standardized data processing protocol for EEG-based analyses of the E/I ratio.

**Highlights:**

- Two EEG spatial filtering schemes - CAR and SL - are compared regarding E/I ratio
- CAR and SL are shown to substantially affect aperiodic spectral slope estimates
- CAR and SL yield opposite conclusions about frontal E/I ratio on the same data
- SL reverses the common *β*_*lo*_ *< β*_*hi*_ bimodal pattern over frontal cortex
- Spatial filtering effects are confirmed on two independent datasets

## 1. Introduction

Optimal information processing in the human brain fundamentally depends on the balance between excitatory (E) and inhibitory (I) neural activity [1]. In line, disruption of the E/I balance is suspected to be involved in a variety of clinical conditions including, but not limited to, autism spectrum disorder [2], schizophrenia [3, 4], Alzheimer’s Disease [5] and other types of dementia [6]. While these observations stress the outstanding importance of the E/I balance, measuring the E/I ratio itself is yet challenging and usually involves invasive techniques and/or interventions [7], thus limiting its clinical utility. It has been proposed lately, however, that the 1*/f* slope of local field potential (LFP) spectra might be a suitable, non-invasive marker of the E/I ratio [8].

Namely, electrophysiological brain activity — as registered via electroencephalography (EEG), magnetoencephalography (MEG) or electrocorticography (ECoG) — commonly exhibits a 1*/f* -type power spectrum, where spectral power |*A*(*f*) | ^2^ follows a power-law dependency on frequency according to |*A*(*f*) | ^2^ ∝*cf* ^*−β*^ with some constant *c*, and the scale-free (or *fractal*) property is characterized via the spectral exponent *β* [9, 10]. Since *β* is equivalent to the coefficient of an ordinary least squares (OLS) regression model after log-log transforming the spectrum, it is often referred to as the *spectral slope* [11, 12]. According to the E/I ratio-slope hypothesis [8], a steeper slope indicates an increase in inhibitory tone, while the opposite holds for an increase in excitatory input. Indeed, several independent works provided support for this phenomenon via pharmaceutical interventions [13, 14, 15], optogenetics [16] or computational modeling [17]. This, combined with recent methodological advances for estimating the 1*/f* slope from empirical data [12, 18] sprouted an immense interest in this field.

While technical developments for obtaining *β* are essential and commended, less emphasis is put on how the data is prepared for slope estimation. This can be a crucial aspect for the broad clinical translatability of the E/I ratio, also acknowledged by Gao and colleagues in their original work as *“slope-inferred E:I ratio should only be interpreted in the context of a comparative experimental design in which the relative E:I ratio can be interrogated in response to experimental manipulations or population differences”* [8]. Indeed, there is significant diversity in the pre-processing pipelines utilized in contemporary literature despite known plausible effects on spectral characteristics [19], limiting comparability of independent outcomes. Another source of variability concerns the frequency range of interest from where *β* is extracted. The association between E/I-ratio and *β* was originally proposed for a fairly narrow range about 20-50 Hz, yet the range of interest varies greatly among recent studies. This aspect is especially important in light of previous evidence implying that the relationship between cortical inhibitory tone and the spectral slope in the low-frequency (i.e., below 5 Hz) regime might be the opposite to that above ∼ 20 Hz [20, 13].

Given the immense potential clinical utility of a reliable E/I ratio index and the considerable research efforts of late, these concerns warrant attention and need to be addressed. In this study we take the first steps towards filling this knowledge gap by explicitly and robustly demonstrating that utilizing alternative spatial filtering techniques — a common step in EEG processing — can lead to conflicting conclusions regarding the E/I ratio even in a confined, comparative experimental environment, while we also confirm the same regarding varying the range of slope estimation.

## 2 Results

We considered the effect of two spatial filtering techniques on estimates of *β*, namely, common average referencing (CAR) and small Laplacian (SL) filtering [21], as they are widely utilized in the EEG field and their effect on 1*/f* spectra can be intuitively understood based on mathematical principles. We employed a trivial, physiological intervention to experimentally modulate the cortical E/I ratio by transitioning from eyes-open (EO) to eyes-closed (EC) resting-state. Closing the eyes evokes a generalized increase in alpha synchronization, which is the most prominent over the parieto-occipital cortices[22] and is often considered as a proxy for increased inhibitory tone [23, 20]. Therefore, we expected a reduction in the E/I ratio in EC compared to EO, more prominently over occipital in contrast to frontal brain areas. The study protocol is illustrated on **Figure 1A** and described in detail in **Materials and Methods**. Importantly, we analyzed two independent EEG datasets with differing electrode montages (**Figure 1B**) to robustly test our hypotheses under multiple experimental conditions. Below we report analyses for Study Cohort #1, while those for Study Cohort #2 are detailed in the **Supplementary Information** and in the main text we limit our discussion to highlight differences in outcomes between Cohort #1 and #2.

**Figure 1:**
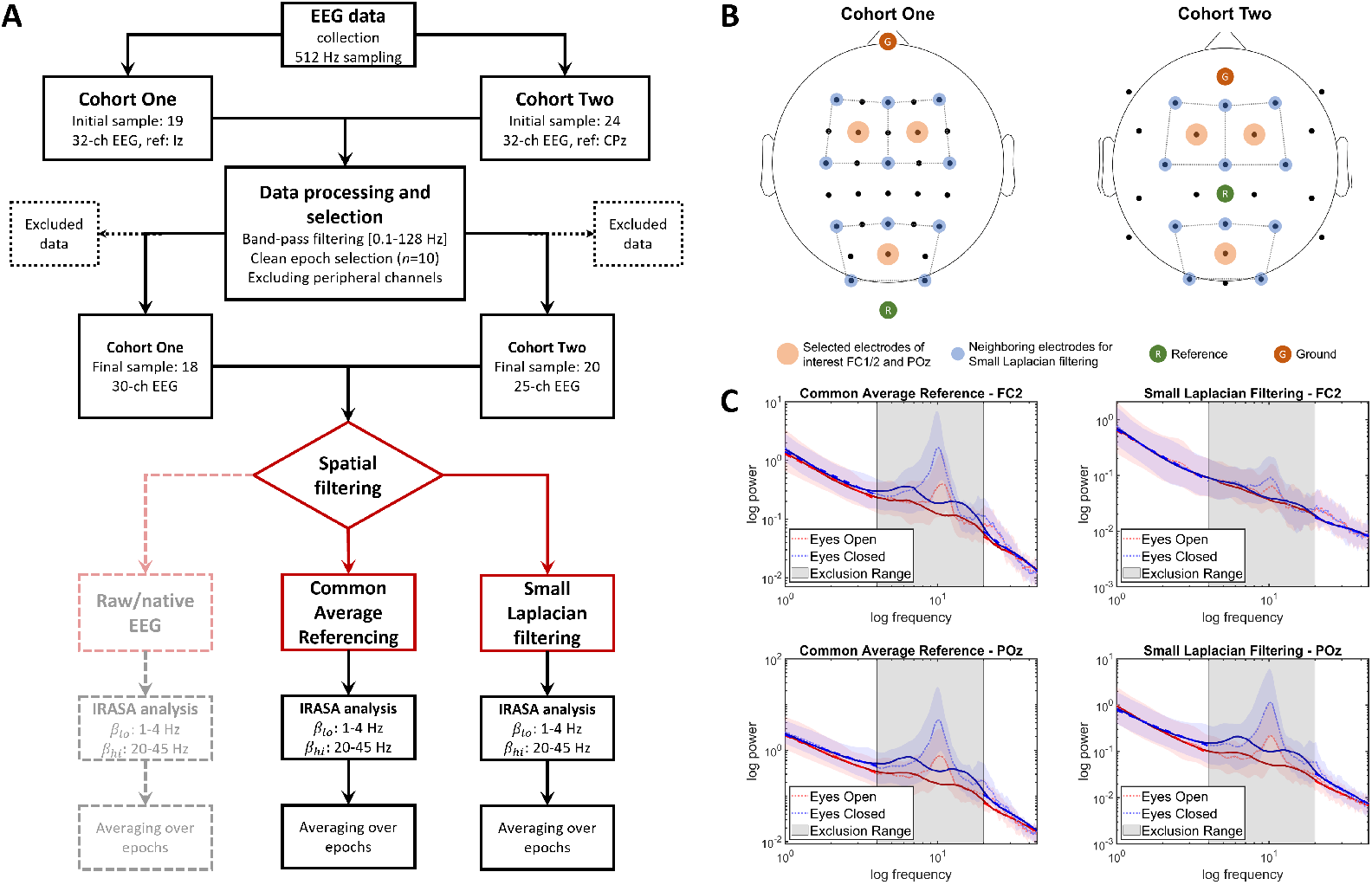
Analysis pipeline. A: Flowchart of analysis steps. We analyzed two independent datasets - Cohort One and Cohort Two - collected via using different EEG electrode montages. The distinctive step in analytical pipelines - spatial filtering - is indicated in red. Besides CAR and SL spatial filtering, we executed the analysis pipeline on raw (i.e., no spatial filtering) data as well. B: The utilized electrode montages. Electrodes of interest are denoted in orange, while the neighborhood utilized for SL filtering is denoted by blue. The selection of representative frontal and posterior cortical regions was guided by practical considerations as so we can perform SL filtering in the two cohorts equivalently. Brown and Green represents the ground and reference electrode positions, respectively. C: Illustration of the IRASA method via group-averaged power spectra obtained via CAR- (left panels) and SL-treated (right panels) EEG data over FC2 (upper row) and POz (lower row). Dotted and continuous lines denote the raw/native and isolated fractal spectra, respectively, while shaded, colored areas mark the corresponding standard error of the mean. The gray area indicates the frequency range excluded from slope estimation (4 − 20 Hz). CAR: Common Average Reference; SL: Small Laplacian; IRASA: Irregular Resampling Auto-Spectral Analysis.

### 2.1 Spectral Slope Analysis - Inferring on the E/I ratio

We selected two representative cortical regions, FC2 (frontal) and POz (posterior); according to our hypothesis, we expected neural correlates to indicate a reduction in E/I ratio over POz. We utilized irregular resampling auto-spectral analysis (IRASA) proposed by Wen & Liu [12] to estimate spectral slopes from isolated fractal spectra, i.e., without the biasing effect of oscillatory signal components. Considering previous literature on EEG spectra bimodality — i.e., the spectrum can be divided into multiple frequency regimes with different scaling exponents [24, 25, 13, 26] —, we first analyzed spectral slopes in the low- (*β*_*lo*_, 1− 4 Hz) and high-frequency regimes (*β*_*hi*_, 20 − 45 Hz) separately in EO and EC (**Figure 1C**).

First, a 4-way repeated measures linear model was evaluated with within-group main effects i) spatial filtering (CAR vs. SL, *filt*), frequency regime (low-vs. high-range, *range*), physiological condition (EO vs. EC, *cond*) and cortical location (FC2 vs. POz, *ch*). As the four-way *filt*× *range* ×*cond*× *ch* interaction was significant (*d* = 17, *F* = 9.7464, *p* = 0.0062, Greenhouse-Geisser corrected) we subsequently evaluated the two processing pipelines separately in 3-way models. The detailed report of these statistical analyses are found in the **Supplementary Tables S1-S3**. The *range* ×*cond* ×*ch* interaction was significant for both CAR (*d* = 17, *F* = 19.857, *p* = 0.0003, Greenhouse-Geisser corrected) and SL (*d* = 17, *F* = 27.852, *p <* 10^*−*4^, Greenhouse-Geisser corrected), and therefore we further broke down the models along *ch*, and the four cases were evaluated via 2-way repeated measures models considering the *range* and *cond* main effects. This evaluation scheme allowed us to test if i) spectra were bimodal (*β*_*lo*_ vs. *β*_*hi*_), if ii) closing the eyes had an effect on *β* estimates, and iii) if this effect differed in low- and high-frequency regimes (*range × cond*). Additionally, we could address our original question and assess if we arrived to the same conclusions with the different processing pipelines. **Figure 2** summarizes results from these analyses; left and right columns show outcomes obtained from the CAR and SL pipelines, respectively, with top and bottom rows illustrating FC2 and POz.

**Figure 2:**
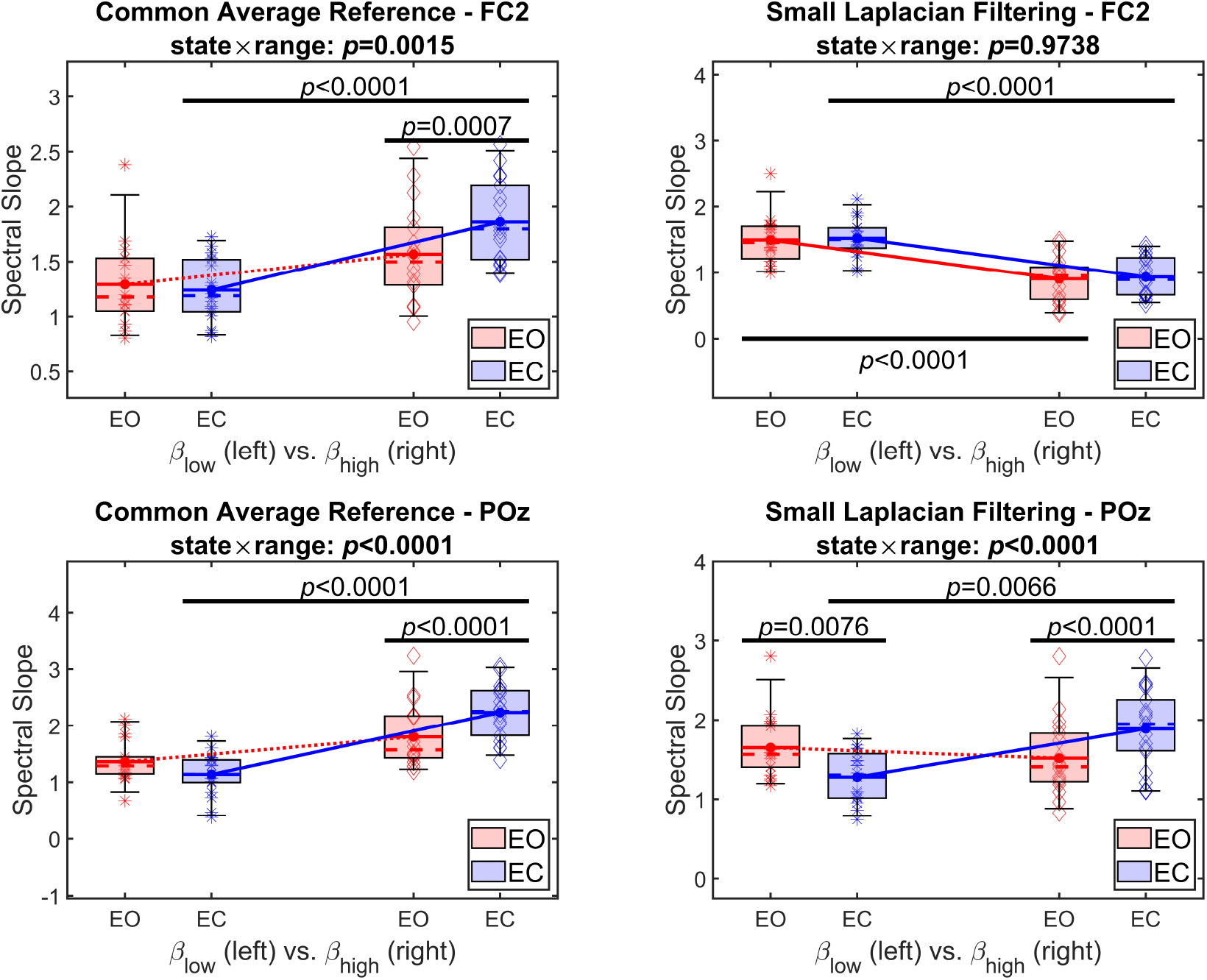
Summary of results from the spectral slope analysis. Left and right panels show outcomes from CAR and SL pipelines, while upper and lower rows depict results from FC2 and POz, respectively. Box plots for *β* estimates from EO are denoted in red, while those from EC in blue. On every panel, box plots on the left illustrate *β*_*lo*_, while those on the right *β*_*hi*_. Significant pairwise differences are denoted by black vertical bars, and the *p*-value for the *state* ×*range* interaction effect is indicated in the panel title. CAR: Common Average Reference; SL: Small Laplacian; EO: Eyes Open; EC: Eyes Closed.

The CAR pipeline exhibited a consistent pattern over FC2 and POz: for both locations, *β*_*hi*_ was significantly higher (i.e., the spectrum was steeper) compared to *β*_*lo*_ (main effect of *range*: *d* = 17, *F* = 16.05, *p* = 0.0009 for FC2 and *d* = 17, *F* = 23.812, *p* = 0.0001 for POz). Post hoc pairwise analyses revealed that this was attributable only to EC (paired t-test, *t*_17_ *<−* 1.9715, *p <* 10^*−*4^ and *d >* 1.8375 for both FC2 and POz), while spectral bimodality could not be confirmed for EO over either location (*p >* 0.1). Finally, *β*_*hi*_ significantly increased over both FC2 (*t*_17_ =− 4.7693, *p* = 0.0007, *d* = 0.7139) and POz (*t*_17_ = − 5.9983, *p <* 10^*−*4^, *d* = 0.8174) in response to closing the eyes, indicating a decrease in E/I ratio over both frontal and posterior brain regions.

On the other hand, SL pipeline produced markedly different outcomes. Over FC2 we only found a significant main effect of *range* (*d* = 17, *F* = 43.71, *p <* 10^*−*5^); however, post hoc pairwise analysis revealed that indeed *β*_*hi*_ was smaller compared to *β*_*lo*_ (paired t-tests, *t*_17_ *>* 6.1480, *p <* 10^*−*4^ and *d >* 1.655 for both EO and EC). Closing the eyes had no significant effect on either *β*_*hi*_ or *β*_*lo*_, suggesting that it did not alter the E/I balance substantially over the prefrontal region of interest. In contrast, a significant *state ×range* interaction (*d* = 17, *F* = 34.882, *p <* 10^*−*4^) over POz warranted detailed pairwise evaluation. This revealed that while closing the eyes decreased *β*_*lo*_ (*t*_17_ = 3.6692, *p* = 0.0076, *d* = 0.9881), it simultaneously increased *β*_*hi*_ (*t*_17_ = −5.771, *p <* 10^*−*4^, *d* = 0.7476); both indicating reduction of the E/I ratio. Additionally, spectral bimodality was confirmed for EC with *β*_*lo*_ *< β*_*hi*_ (*t*_17_ = −3.7358, *p* = 0.0066, *d* = 1.4231), while there was no statistically significant difference between low- and high-range spectral slopes in EO.

In summary, these results indicate that the pre-processing pipeline can affect study conclusions regarding the E/I ratio: while the pipeline utilizing CAR filtering would imply that the E/I ratio decreased over both frontal and posterior regions in response to closing the eyes, the SL pipeline in turn shows that this effect only takes place over POz but not FC2. In addition, while *β*_*lo*_ *< β*_*hi*_ bimodality was observed over both locations in EC for CAR, applying SL reversed this pattern over FC2 to *β*_*lo*_ *> β*_*hi*_.

### 2.2 Resting Alpha Power - The neural proxy for inhibition

To confirm the expected effect of closing the eyes we computed isolated oscillatory alpha band limited power 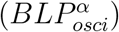 by subtracting log-transformed fractal power from log-transformed total (mixed) power and integrating it over the 8 − 12 Hz range. While 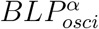 was significantly higher in EC compared to EO over both FC2 (*t*_17_ = −5.1803, *p* = 0.0003, *d* = 1.0723) and POz (*t*_17_ = −4.0277, *p* = 0.0035, *d* = 1.1235) for CAR-filtered data, this difference was only observed over POz (*t*_17_ = −3.1962, *p* = 0.0212, *d* = 0.9067) in the SL pipeline (**Supplementary Figure S1**). These outcomes are in line with expectations on the physiological effect of transitioning from EO to EC.

We also wanted to investigate how the observed changes in *β* relate to the changes in our neural proxy of inhibitory tone, alpha power. To this end we computed difference indices 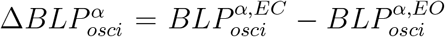 and 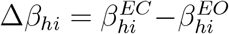 (and equivalently for *β*_*lo*_), where the subscript *osci* denotes that isolated oscillatory BLP was used, while superscript *EO* or *EC* indicates the physiological state. Results of this analysis are depicted on **Figure 3**.

**Figure 3:**
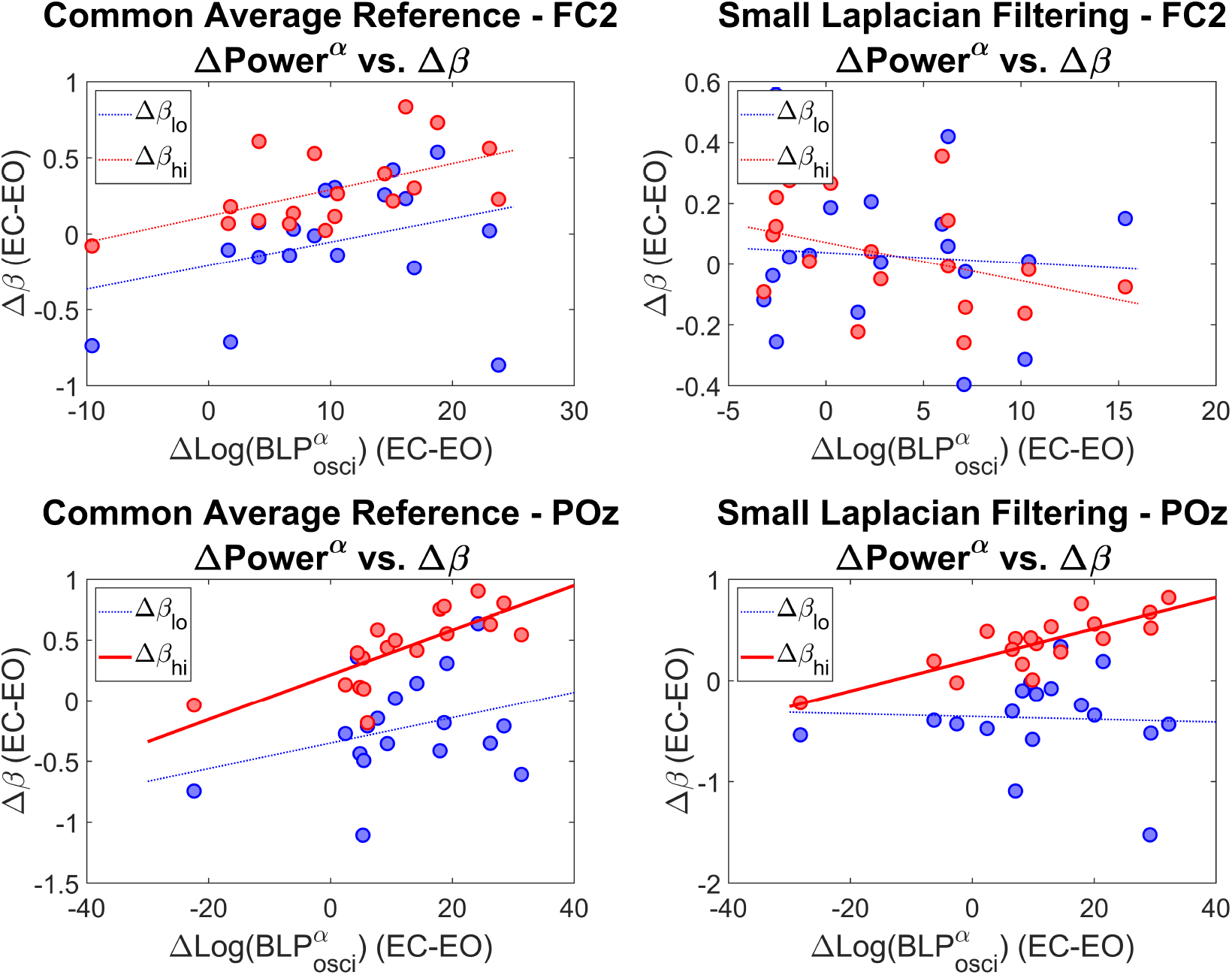
Correlation between the physiological state-related change in isolated oscillatory alpha power and spectral slope. Left and right panels show outcomes from CAR and SL pipelines, while upper and lower rows depict results from FC2 and POz, respectively. Blue dots denote change in *β*_*lo*_ in response to closing the eyes obtained as 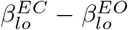, while red dots those in *β*_*hi*_ equivalently. Straight lines illustrate the ordinary least squares fit on samples of corresponding color; thick continuous lines indicate significant correlation between ∆BLP and ∆*β* (Pearson or Spearman depending on data normality), while dotted lines indicate otherwise. CAR: Common Average Reference; SL: Small Laplacian; BLP: Band Limited Power.

In that we found that ∆*β*_*hi*_ and 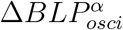 was significantly correlated only over POz in both CAR (Pearson *ρ* = .7540, *p* = 0.0.0024) and SL (Pearson *ρ* = .8117, *p* = 0.0003) pipelines. Notably, no correlations were identified between ∆*β*_*lo*_ and 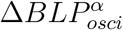

### 2.3 Confirmatory Analyses - Hemispheric laterality

To avoid our outcomes to be biased by potential hemispheric laterality, we executed our entire analytical pipeline using FC1 instead of FC2, too. Overall, outcomes of this analysis are well in line with those found for FC2 (**Supplementary Figure S2**). While there was a significant *state ×range* interaction over FC1 (*d* = 17, *F* = 11.482, *p* = 0.0035) when utilizing SL, post-hoc pairwise comparisons revealed no statistically significant state-related changes in *β*_*lo*_ or *β*_*hi*_.

Notably, we carried out our analytical pipeline for regions CP1 and CP2, the other two EEG channels were SL filtering could be applied equivalently in Study Cohorts #1 and #2 (see **Figure 1B**). These results are depicted on **Supplementary Figure S3** and they show that while the CAR pipeline indicated an increase in *β*_*hi*_ in response to closing the eyes over both CP1 (*t*_17_ = −4.5624, *p* = 0.0011, *d* = 0.5279) and CP2 (*t*_17_ = −5.4754, *p* = 0.0002, *d* = 0.4808), no significant state-related changes in *β*_*hi*_ could be detected over either cortical location when EEG was spatially filtered with SL. In addition, the reversal of the *β*_*lo*_ *< β*_*hi*_ pattern due to SL filtering can also be observed over both CP1 and CP2. These outcomes further confirm that applying an alternative processing pipeline can influence the inferences made on cortical E/I ratio.

### 2.4 Broadband unimodal spectral slope analysis

In order to assess the relevance of treating the spectrum as bimodal, we also analyzed the effect of closing the eyes on spectral slopes obtained unimodally from a broadband 1 −45 Hz range (*β*_*bb*_). Note that such a broadband approach is also common in the literature (see e.g., [14, 16, 17]). Results of this analysis are shown on **Supplementary Figure S4**. As we no longer considered the effect of *range*, we employed 2-way models with within-group effects *state* and *ch* independently in the CAR and SL pipelines. The two cases yielded coinciding results: there was no statistically significant response to closing the eyes over any cortical regions (Fc1, Fc2, POz) in either pipelines, Note that when only considering POz, *β*_*bb*_ was even significantly decreased in both the CAR (*t*_17_ = 2.1856, *p* = 0.0431 unadjusted, *d* = 0.3914) and SL (*t*_17_ = 2.5033, *p* = 0.0228 unadjusted, *d* = 0.5356) pipelines. Therefore, estimating *β* from a broadband spectra might even lead us to conclude that closing the eyes increased the E/I ratio, even though it is contradictory to our neurophysiological expectations. While correlation analysis did not indicate a significant relationship between the change in *β* and oscillatory alpha BLP for CAR, a negative correlation between ∆*β*_*bb*_ and 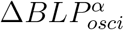 was significant (Pearson *ρ* = −.6718, *p* = 0.0091) over POz in the SL pipeline.

### 2.5 Independent outcomes from Study Cohort #2

Finally, since the reference scheme can have a significant effect on spectral characteristics [19], we wanted to ensure that our findings are not specific to the recording setup and thus we carried out our entire analysis pipeline on an independent set of recordings. In general, outcomes obtained from Cohort #2 were comparable to those of Cohort #1 with some notable exceptions (see **Supplementary Tables S4-S6** for details on 4- and 3-way models). Bimodal slope analysis over FC2 and POz (**Supplementary Figure S5**) were similar as shown in **Figure 2**, only we did not observe a decrease in *β*_*lo*_ over POz in response to closing the eyes. Similarly to Cohort #1, results for FC2 and FC1 were identical in nature (**Supplementary Figure S6**). On the other hand, in Cohort #2 the CAR and SL pipelines yielded coinciding results in case of CP1 and CP2, with both spatial filtering schemes indicating an increase in *β*_*hi*_ when closing the eyes for both locations (**Supplementary Figure S7**). As another notable difference from Cohort #1, broadband analysis identified a strong, state-related decrease in *β*_*bb*_ over POz on both pipelines (**Supplementary Figure S8**), indicating that closing the eyes resulted in a reduction of inhibitory tone over POz. While the expected changes in alpha BLP were replicated (**Supplementary Figure S9**), no correlations were found between 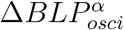 and ∆*β*_*hi*_, ∆*β*_*lo*_ or ∆*β*_*bb*_. In summary, these analyses further confirmed that the choice of spatial filtering technique, as well as that of the scaling range for slope analysis can fundamentally affect study conclusions, and these effects were consistent regardless of electrode montage used for data collection.

## 3 Discussion

### 3.1 The bimodal approach and the influence of spatial filtering

The multimodal nature in the power spectra of electrophysiological recordings is well established [24, 25, 27, 26], and previous literature indicates that it can have a seminal effect on how estimates are interpreted in the context of E/I ratio. Becker and colleagues [20] employing resting alpha power as a proxy for inhibitory tone based on the “Gating by inhibition” hypothesis [23] reported that the power spectrum becomes flatter (reduced *β*) in the low-frequency regime (*<* 5 Hz) with the increase of E/I ratio. More importantly, Muthukumaraswamy and Liley [13] showed that after administering Tiagabine — a GABA agonist increasing inhibitory tone —, a simultaneous decrease in low-range slope *β*_*lo*_ (in line with [20]) and increase in high-range slope *β*_*hi*_ (in line with [8]) could be observed. Despite these reports, the bimodal nature of the power spectrum is most often neglected in recent works probing the E/I ratio via the spectral slope. While numerous recent studies limited their range for estimating *β* roughly to the 30 − 50 Hz [15] in line with [8], others considered frequency regimes spanning approximately 2− 25 Hz [28, 29, 30, 31] or broadband 1− 45 Hz [16, 14, 17], and only a handful of articles reported analysis outcomes treating low- and high-range slopes separately [13, 26, 32]. Given our results here we argue that spectral slope estimation should be implemented separately for low- and high-frequency regimes, not solely based on the fact that *β*_*lo*_ and *β*_*hi*_ estimates can be statistically significantly different, but because they apparently respond to change in physiological state in varying ways [13]. We too observed this effect when analyizing SL-fitlered data over POz (**Figure 2**), but more importantly, while our bimodal analyses consistently pointed towards a decrease in E/I ratio due to closing the eyes, the broadband unimodal analysis would indicate the opposite (**Figs. S4** and **S8**).

Notably, there is substantial variability in data processing pipelines among previous literature. CAR is a frequently applied spatial filtering step [13, 16, 31], while re-referencing to mastoids [29] or earlobes [30] is also common. Additionally, many studies employ more elaborate manipulations to the data such as independent component analysis (ICA)-based artifact removal [29, 13, 14, 31, 15] or source activity reconstruction [15, 17], which could introduce further effects on slope estimates that are beyond the scope of this work. This is not only relevant in terms of the E/I ratio hypothesis but also bimodality, as our results showed that the previously reported pattern of *β*_*lo*_ *< β*_*hi*_ [24, 25] can not only be affected, but even reversed by the selected spatial filtering technique (e.g., **Figure 2** or **Supplementary Figure S5**). These observations call for a standardized data processing — and likely, analytical — pipeline for establishing consistency and comparability among independent works.

We also wanted to benchmark our outcomes against those obtained from raw EEG data i.e., in native electrode space with no spatial filtering applied. While our main results were mostly confluent among the two study cohorts, distinctive characteristics emerged from raw data that point towards the influence of the reference position. The previously observed bimodality pattern of *β*_*lo*_ *< β*_*hi*_ was ubiquitous for Cohort #2, while present only in EC in Cohort #1 (over all cortical locations). In Cohort #1 (**Supplementary Figure S10**), results from raw EEG reflected those from the CAR pipeline over FC1 and FC2, however those from the SL pipeline over POz, with a simultaneous state-related decrease in *β*_*lo*_ (*t*_17_ = 4.0001, *p* = 0.0037, *d* = 0.8654) and increase in *β*_*hi*_ (*t*_17_ = − 5.3808, *p* = 0.0002, *d* = 0.6687). On the other hand, in Cohort #2 (**Supplementary Figure S11**) raw- and CAR-derived outcomes were almost identical. This further highlights the importance of the reference electrode position regarding spectral EEG features (also see below), while at the same time implies that simple re-referencing of EEG channels [29, 30] can influence slope estimates in unexpected ways.

### 3.2 Insights on the relationship between spectral slope and E/I ratio

We followed the same assumption as Becker and colleagues [20] — also supported by recent literature [33, 34] — in taking resting alpha power as a proxy for inhibitory tone [23]. Consequently, we hypothesized that closing the eyes will produce a stronger increase in inhibition over occipital compared to frontal brain areas [22], which was supported by alpha BLP analysis (**Supplementary Figure S1**). In this framework, our results suggest that SL filtering might be superior to CAR with regards to drawing inferences on the E/I ratio. Precisely, from CAR-treated data one would conclude that the E/I ratio decreased due to closing the eyes over both frontal (FC1, FC2) and posterior (POz, CP1, CP2) areas as indicated by an increase in *β*_*hi*_, while from EEG data spatially filtered with SL, one would came to this conclusion only with regards to POz in Cohort #1 (**Supplementary Figure 2**), and POz, Cp1 and Cp2 in Cohort #2 (**Supplementary Figures S5-S7**). Furthermore, only with SL spatial filtering (or analyzing raw data) could we observe a simultaneous decrease in *β*_*lo*_ and increase in *β*_*hi*_ when transitioning from EO to EC, in line with Becker et al. [20] and Muthukumaraswamy & Liley [13].

It is worth noting that while results obtained from the SL pipeline are closely comparable between Cohorts #1 and #2, they are not identical despite the full equivalence of the spatial filtering technique (unlike for CAR, where the number of channels and their arrangement were different in the two cohorts). This provides further support to the — expected [19] — influence of the reference position: In Cohort #2 the reference (CPz) was approximately the same distance from the two regions of interest, while in Cohort #1 (Iz) it was in close proximity to POz but not to FC2/FC1 (**Figure 1**). It is well established that the reference position can have significant influence on EEG signal power [35] or estimated brain network structure [36], but less is known about its potential confounding effects on the spectral slope [37]. Although we did not explore this aspect any further here, this appears as an interesting question that we intend to pursue in the future.

We found a significant positive correlation between 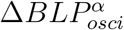 and ∆*β*_*hi*_ over POz (**Figure 3**). By taking alpha BLP as a proxy for inhibitory tone these results can indicate that the observed increase in *β*_*hi*_ is related to increased inhibitory tone; however, this also raises the concern that resting alpha power might confound slope estimates, especially in the high-frequency regime. Addressing this issue in the current setup is challenging, given that the two neural indices — alpha BLP and *β* — might capture the same phenomenon, and thus their correlated nature might stem from neurophysiological and not methodological reasons. Recent evidence is supporting this, where alpha bursts with high amplitude were suggested to represent short periods with stronger inhibition, alternating with excitation/-disinhibition [33, 34]. Further elucidating this question also appears to be possible via experimental modulation of the E/I ratio (see **Limitations and future work** below).

Nevertheless, we tentatively draw two overall conclusions from these ramifications: foremost, it is robustly confirmed that CAR and SL filtering could lead us to conflicting inferences on the change of E/I ratio in response to closing the eyes, as well as SL — and other spatial filtering techniques promoting localization of neural activity, such as current source density (CSD) transformation [38] — appears as the suitable choice for data processing when one sets out to analyze the spectral exponent of EEG data.

### 3.3 Why spatial filtering might affect spectral slope estimates

The results presented here empirically highlight the seminal importance of the selection of spatial filtering technique in EEG spectral slope analysis; however, they do not allow insight on the origins of the observed phenomena. While we believe that a systematic investigation of this question is beyond the scope of this study, below we make an attempt to address this question from an intuitive angle.

While CAR and SL serve different purposes and therefore both have their preferred use cases [21], they share a common feature: both involve the summation (averaging) of multiple EEG time series, that are then subtracted from individual channel data. Importantly, this manipulation does not simply involve addition of two (or more) independent, stochastic time series but instead those with a 1*/f* spectra. The superposition of two 1*/f*^*β*^ processes with different spectral exponents *β*_1_ and *β*_2_ can introduce a crossover in the power spectrum — or equivalently, the scaling function in the time domain — of the composite time series [39, 40, 41, 27]: given the superposition of two theoretical processes with infinite length and sampling and with *β*_1_ *> β*_2_, the variance profile of process 2 will eventually dominate that of process 1 as frequency increases, resulting in a power spectrum with a steeper slope (dominated by *β*_1_) for lower frequencies and a slope close to *β*_2_ for high frequencies [40, 41, 42]. An illustration of such a case is presented on **Figure 4A**. Theoretically, this argument holds for the superposition of an arbitrary number of time series (for an example, see **Figure 4B**), as well as applies similarly to processes with known bimodal spectra, regardless if *β*_*lo*_ *< β*_*hi*_ (**Figure 4C**) or *β*_*lo*_ *> β*_*hi*_ (**Figure 4D**).

**Figure 4:**
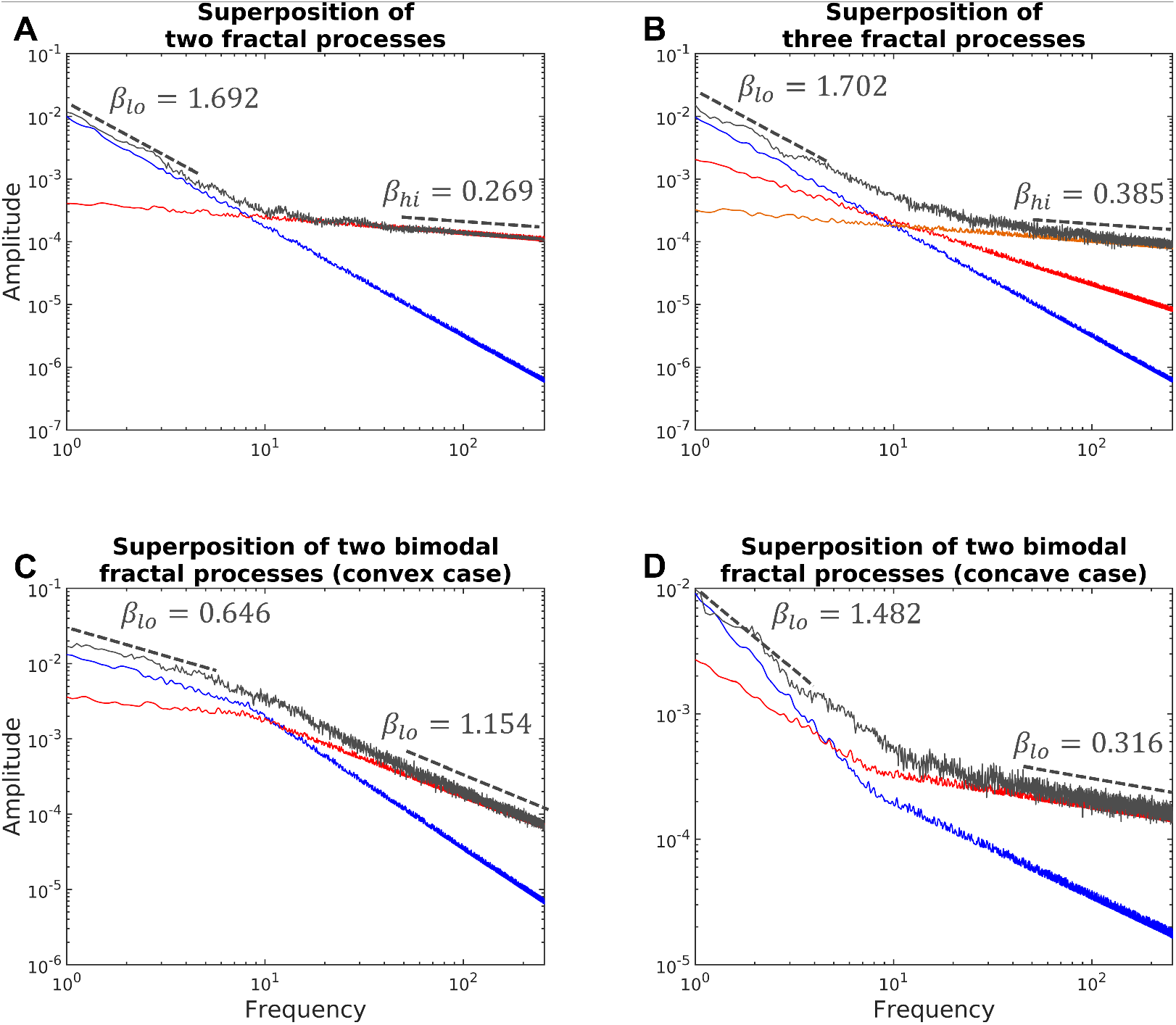
Illustration of how the superposition of multiple 1*/f* signals can result in a crossover in the power spectrum (Gray: spectra of composite time series). A: superposition of two 1*/f* signals with theoretical *β* = 1.75 (blue) and *β* = 0.25 (red). The power spectrum of the composite time series is dominated by the larger exponent in the low-, while by the smaller in the high-frequency regime. B: superposition of three 1*/f* signals with *β* = 1.75 (blue), *β* = 1 (red) and *β* = 0.25 (orange). The spectral slope is dominated by the largest exponent in the low-, while by the smallest in the high-frequency regime. C: superposition of two bimodal signals with *β*_*lo*_ *< β*_*hi*_; for the blue signal, *β*_*lo*_ and *β*_*hi*_ was set to 0.75 and 1.75, respectively, while for the red signal *β*_*lo*_ = 0.25 and *β*_*hi*_ = 1. Apparently, the low-frequency range is dominated by the blue signal with larger exponent in the given range, while the opposite holds for high-frequency range. D: superposition of two bimodal signals where *β*_*lo*_ *> β*_*hi*_; for the blue signal, *β*_*lo*_ and *β*_*hi*_ was set to 1.75 and 0.75, respectively, while for the red signal *β*_*lo*_ = 1 and *β*_*hi*_ = 0.25. The previous observations regarding the dominant spectral exponent still hold.

Nevertheless, these arguments only provide intuitive support but not a sound explanation. Most importantly, the arguments only hold precisely for time series that are independent, which is clearly untrue for EEG channels, as supported not only by the vast literature on EEG-derived functional connectivity studies [43], but also self-evident from the volume conduction effects inherent to EEG methodology [44]. This might explain why we observed the reversal of the *β*_*lo*_ *< β*_*hi*_ over frontal regions only for SL but not for CAR, yet answering this rigorously requires further research. Moreover, they cannot fully capture and formally express the effects of non-trivial manipulations to data such as CSD transformation or ICA-based noise removal on the power spectral slope. For this reason, in this confined study we intentionally limited our exploration to techniques that are easily understandable (namely, CAR and SL). Nevertheless, this question is seminal for the derivation of a standardized data processing pipeline with a sound theoretical foundation and warrants further research; which is our central goal in the future.

### 3.4 The potential influence of the method for obtaining slope estimates

A rightful question can be raised regarding our choice of the evaluation technique, IRASA [12] and its parameter settings. IRASA was published in 2016 — building on concepts first introduced in 1991 [45] —, however another popular method termed ‘fitting fitting oscillations & one over f’ (FOOOF) was introduced recently (in 2020) to separately characterize the broadband 1*/f* and superimposed oscillatory components of neurophysiological signals [18]. FOOOF gained immense popularity in the past years and likely is the most frequently used technique for similar analyses (see e.g., [14, 30, 16, 15, 17]). FOOOF arguably has advantages over IRASA in terms of allowing for a broader range for parametrization of oscillatory components, while the latter puts the emphasis on the unbiased estimation of the spectral slope along theoretical assumptions. Relatedly, while FOOOF implements a data driven, recursive approach to separate the broadband, aperiodic component from narrow-band oscillations, IRASA is a model-based technique assuming an additive signal model and exploiting the theoretical self-affinity property of the presumed fractal signal component. Overall, both methods can have favorable properties in a given situation [18], yet we decided to implement our analysis pipeline employing IRASA for one main reason. It has been observed previously that human neurophysiological recordings can indeed express power spectra where *β*_*lo*_ *> β*_*hi*_ [46, 47, 27]. FOOOF models the aperiodic component as a Lorentzian (instead of a pure power-law) function, allowing for a ‘knee’ parameter [18]. This model inherently allows for the characterization of the slope only in the high-frequency range (i.e., over the ‘knee’) and thus is unsuitable for treating bimodal signals where *β*_*lo*_ *> β*_*hi*_. IRASA does not suffer from this limitation, although it can cause a ‘smoothing’ of the breakpoint in the spectrum depending on its parameter settings [12]. We took this into consideration when designing our analysis pipeline and accordingly excluded the range from slope estimation where the breakpoint was expected. Nevertheless, as a sanity check we executed our analysis pipeline using FOOOF with corresponding parameter settings; results of which can be accessed in detail in the **Online Supplementary Information**. In general, outcomes obtained via FOOOF aligned well with those of IRASA. In Cohort #1, an increase in *β*_*hi*_ in response to closing the eyes were only found over POz regardless of CAR or SL (**Supplementary Figures S12-S13**). A simultaneous decrease in *β*_*lo*_ was observed only over POz with SL. In addition, we did not observe any state-related change in *β*_*bb*_ in any of the cases (**Supplementary Figure S14**) when performing broadband (1− 45 Hz) FOOOF analysis.

In Cohort #2 we observed similar contradictory results over Fc2, in that the CAR pipeline indicated an increase in *β*_*hi*_ over Fc2 but not the SL pipeline (**Supplementary Figure S15**. However, considering Fc1 instead showed an increase in *β*_*hi*_ in both the CAR and SL pipelines (**Supplementary Figure S16**). We also observed a decrease in *β*_*bb*_ due to closing the eyes over FC2 (**Supplementary Figure S17**) in Cohort #2, again, contradictory to what was expected. Overall, these analyses yielded similar conclusions in that differing spatial filtering pipelines can lead to contradictory conclusions on E/I ratio, regardless if the spectral slope is assessed via IRASA or FOOOF. Nevertheless, a combination of IRASA and FOOOF appears reasonable to take advantage of both methods in the future, e.g., to use IRASA to characterize the 1*/f* spectra in low- and high-frequency regimes separately regardless of the nature of bimodality, while utilizing FOOOF to parametrize oscillatory peaks.

To further explore the aspect, we also present preliminary results in the **Online Supplementary Material** from an analysis pipeline where Cohort #1 and #2 datasets were evaluated with a novel technique (*in development*), which we refer to as MultiModal Spectral Parametrization Method (MM-SPM). This technique utilizes the recursive, data-driven approach for separating broadband 1/f from oscillatory components based on the FOOOF algorithm [18]; however, a key difference is that instead of assuming a Lorentzian model for the former, MMSPM employs a piece-wise linear regression framework with the constraint that the broadband, locally 1/f component is continuous (i.e., there is no sudden change in power on the border of different scaling ranges, as illustrated on **Supplementary Figure S18**). While we do not present details of this method here as it is expected to be published elsewhere, preliminary analysis outcomes are as follows (using identical processing and equivalent parameter settings to the IRASA and FOOOF analyses). In Cohort #1, no state-related response in either *β*_*lo*_ or *β*_*hi*_ were observed in the CAR pipeline, while the SL pipeline denoted an increase in *β*_*hi*_ over POz and Fc1. In Cohort #2, both CAR and SL yielded a simultaneous decrease in *β*_*lo*_ and increase in *β*_*hi*_, however while CAR also indicated an increase in *β*_*hi*_ over FC1 and FC2, this was not the case for SL over either location. Very importantly, the MMSPM method also reliably showed that the *β*_*lo*_ *< β*_*hi*_ pattern that is observed over all locations with CAR is reversed over FC1 and FC2 to *β*_*lo*_ *> β*_*hi*_ when using SL. Figures illustrating all these outcomes can be accessed in the **Online Supplementary Material**. In summary, outcomes obtained with this new procedure (still *in development*) confirm two notions: i) application of different spatial filtering schemes can yield contradictory results regarding the E/I ratio, and ii) the spectral slope should be estimated via a technique that can account for cases where *β*_*lo*_ *> β*_*hi*_.

### 3.5 Limitations and future work

Most importantly, our work is empirical in nature i.e., we do not yet provide a full and sound explanation of the observed effects based on theoretical formulation and computer simulations. While we acknowledge this limitation, we believe that the presented results carry significant value in themselves as they raise awareness to an important phenomenon that can fundamentally affect study conclusions and thus lead to controversy in the literature. It is nevertheless one of our immediate future goals to redeem these constraints. Furthermore, in terms of interpretation we could only rely on suppositions in contrast to explicit neuroscientific evidence. Namely, we made two key assumptions based on previous literature: first, we took resting alpha power as a proxy for inhibitory tone [23, 20, 48, 20] and thus EC resting as a state of increased inhibition; and second, we hypothesized that this response in inhibitory tone will be more prominent over occipital compared to prefrontal regions [22]. Therefore, while the outcomes most consistently observed for SL-treated data are aligned with these initial assumptions, we cannot contrast them with ‘ground truth’ measurements of cortical E/I ratio or inhibitory tone in the investigated locations. Although we believe that this does not reduce the technical relevance of the results presented here, it definitely emphasizes the need for more elaborate investigation where the E/I ratio is explicitly and selectively controlled with some appropriate neuromodulation technique such as transcranial electrical [49] or magnetic [50] stimulation, and ground truths are obtained e.g., via sensory evoked potentials [51]. Therefore, it is also amidst our immediate goals to replicate our results in such an environment.

Besides these notions, we can identify additional aspects where we can extend upon our work. Our analyses were constrained by technical details, as we had to select POz and FC2 (FC1) as regions of interest due to the available electrode montages (**Figure 1**). Higher spatial resolution EEG would allow us to more specifically select brain regions suitable for a given employed intervention paradigm. Also, we did not employ a personalized approach for assessing alpha activity around the individual peak frequency [52] but instead characterized alpha power universally in the 8 −12 Hz range. FOOOF could indeed enhance this aspect for our investigation by parametrizing alpha peaks — and thus help fine tuning low- and high scaling ranges, too — for each participant individually [18]. The relationship between the change in spectral exponents and resting alpha power was only assessed via correlation analysis due to the characteristics of the data, however this does not necessarily prove causality i.e., that a change in E/I ratio was the reason for the change in the spectral slope. Furthermore, this relationship was only significant for *β*_*hi*_ but not *β*_*lo*_ in any of the cases, which calls for more caution when linking *β*_*lo*_ to E/I ratio despite previous reports [20, 13]. Finally, our analysis here was limited to EEG, though same considerations should be made for analyzing the 1*/f* properties of neural activity recorded with other modalities such as magnetoencephalography [53, 15].

### 3.6 Summary

To summarize, in this study we provided empirical evidence that the choice of pre-processing pipeline can lead to outcomes with conflicting interpretations even in an isolated, comparative experimental environment. We confirmed this notion robustly on two independent datasets, while also pointing out limitations and potential directions for future research. Overall, our results suggest that a spatial filtering step that aims on improving spatial resolution and localization of neural activity should be preferred when estimating the spectral slope. Given the growing interest surrounding cortical E/I ratio and how it might be affected in various physiological and pathological conditions, our results call for the establishing of a standardized evaluation protocol, so to render studies with similar interest as confluent and comparable as possible.

## 4 Materials and Methods

### 4.1 Participants and data collection

EEG data was collected in overall three independent studies with non-overlapping cohorts. While study protocols differed, they had the same inclusion/exclusion criteria (see below) and resting-state EEG was collected at the very beginning of every recording session, thus rendering the datasets comparable. All projects were conducted in line with the regulations of the Declaration of Helsinki, and the protocols were reviewed and approved by the Institutional Review Board of The University of Texas at Austin (approval number: 2020-03-0073). All participants provided written informed consent before the measurement, and recordings were carried out in an isolated room with dim lighting, while participants were sitting in an armchair. All participants were aged between 18 and 40 years, had no reported history of neuropsychiatric or other major medical condition (e.g., open heart surgery), and were free of medication affecting brain function.

The first study cohort (Cohort #1) consisted of 19 young, healthy volunteers (mean age 27.26 ± 6.11 years, 10 females, all right handed). EEG data was collected using an ANT neuro EEGO amplifier and a customized cap that monitored neural activity from 32 of the international 10-5 standard locations with high electrode density over the mid- and parasagittal lines (**Figure 1B**, left). The reference electrode positioned over the inion (Iz). Note that channels AF3 and AF4 were excluded from analysis due to their susceptibility to eye movement-related components, yielding 30-channel EEG data. The recording protocol involved two minutes of EEG monitoring with eyes open followed by two minutes with eyes closed, with a 10-second intermediary period in between. EEG data in the second study cohort (Cohort #2) was collected in two independent studies utilizing the same equipment. The total sample consisted of 24 young, healthy individuals (mean age 22.96± 2.63 years, 7 females, 22 right handed). EEG data was collected using a similar ANT neuro EEGO amplifier from 32 (international 10-10 standard) scalp locations, with the reference electrode positioned over CPz (**Figure 1B**, right). Note that channels FP1, FPz, FP2, T7, T8, M1 and M2 were excluded from analysis as these channels are more commonly affected by eye movements, blinks, and skeletal muscle artifacts, yielding 25-channel EEG data. Data collection protocol was similar to that of Cohort #1, only in these recordings either one or two minutes of EO and EC data was collected.

In both cohorts, EEG data was sampled at 512 Hz, and all electrode impedances were kept under 5 kΩ. During both conditions participants were instructed to rest, not to engage in any specific mental activities, refrain from motion and stay awake. During EO recordings, a fixation cross was presented on a computer screen, on which participants were asked to focus their gaze on to minimize eye movements.

### 4.2 Data pre-processing and selection

All data analyses, including pre-processing, obtaining of neural indices and statistics, were implemented and carried out in Matlab 2023b (The MathWorks, Nattick, USA). The data analysis pipeline is illustrated on **Figure 1A**. Raw data was band-pass filtered using a 4^th^-order, zero-phase Butterworth filter with cutoff frequencies 0.1 and 128 Hz. Then, segments containing absolute amplitude values above 100 *µ*V in any of the EEG channels were excluded with additional padding of 1/16 s. Finally, segments with a length of 4 seconds were selected randomly from the remaining data with a maximal overlap of 3.5 s, and those participants where this procedure could not yield at least 10 clean epochs in both conditions were also excluded from further analysis. This resulted in a final sample size of 18 and 20 participants in Cohorts #1 and #2, respectively. Finally, data epochs were subjected to one of the two spatial filtering schemes as detailed below.

As a representative frontal/prefrontal channel, we selected the FC2/FC1 location, while POz as a representative occipital/parietooccipital location. The reasoning for these picks stems from the electrode montage utilized in the two cohorts (see **Figure 1B**); namely, these were the most frontal and occipital available scalp locations for which we could apply SL filtering in the same exact manner among the two cohorts (see below).

Data was either re-referenced to the common average electrode (CAR scheme) or subjected to small Laplacian filtering (SL scheme). During CAR, the average EEG signal computed over all channels and then subtracted from each individual channel. In the SL scheme, average activity of neighbouring channels was computed and then subtracted from each respective channel [54]. For FC2 (and equivalently, FC1) these included locations Fz, F4 (F3), Cz and C4 (C3), while P3, Pz, P4, O1 and O2 for POz.

### 4.3 Spectral slope analysis

Spectral slope estimates were obtained using the IRASA technique introduced by Wen and Liu [12]. This method assumes a composite model where neural activity is treated as a superposition of scale-free/fractal and oscillatory/periodic components. In this framework one can exploit the self-affine property of broadband scale-free signals [55], in that their power spectral characteristics are unaffected in distribution by resampling — and only subject to rescaling —, while this property does not hold for harmonic signals. Along these principles, one can devise a method to isolate the power spectra corresponding to the fractal component by taking the geometric mean of the power spectra obtained from up- and downsampled versions of the original signal [45, 56]. IRASA improves upon this principle by utilizing a set of non-integer resampling factors thus to reduce the probability of retained oscillatory peaks due to harmonic overlap [12]. After the scale-free spectrum is isolated, the spectral exponent (slope) can be estimated via OLS regression of log-transformed power over log-transformed frequency without the bias of present harmonic components such as alpha or theta peaks, or line noise and its harmonics at 60 and 120 Hz [12]. In addition, the isolated spectra of oscillatory components can also be obtained by subtracting the scale-free from the original power spectrum, assuming no phase coupling between scale-free and oscillatory components. For more details on the IRASA technique and its precise algorithm, please refer to [12] or [57].

IRASA was employed with the following parameter settings in the two processing pipelines (CAR and SL). The set of resampling factors was set to default ranging from 1.1 to 1.9 in increments of 0.05, as recommended by Wen and Liu [12]. Spectral slopes were estimated from the fractal spectra in the 1 −4 Hz (low-) and in the 20− 45 Hz (high-) ranges. The boundary frequencies were set so the filtering effects introduced by resampling, as well as the documented ‘smearing’ of the spectra around the prominent alpha peak do not intersect with the ranges of interest (for more details see the Supplementary Material in [26]). To obtain robust estimates of the EEG power spectrum, IRASA was carried out for each 4-second epoch, and then corresponding spectra were averaged over all windows before obtaining spectral slopes.

Spectral exponents for the low- (*β*_*lo*_) and high- (*β*_*hi*_) ranges were obtained from these averaged spectra. Additionally, to characterize inhibitory tone, we estimated log-transformed oscillatory alpha band limited power (BLP). In that the proportion of oscillatory power was computed by subtracting the log-transformed scale-free spectrum from the log-transformed raw power spectrum, and then summed over the 8 − 12 Hz range 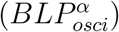. To measure the effect of closing the eyes on spectral features, difference measures were obtained as 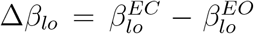 (and equivalently for *β*_*hi*_) and 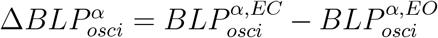 where EO and EC in superscript denotes the physiological condition. Note that even though EEG spectral exponents are generally negative i.e., power decreases as frequency increases, conventionally we refer to *β*_*lo*_ and *β*_*hi*_ through their absolute values [11] and thus a larger *β* value indicates a steeper spectrum/slope and vice versa.

### 4.4 Statistical analyses

All variables were first tested for normality using the Lilliefors test. Since all outcome measures regarding the spectral slope satisfied the normality criterion, data was then first evaluated via a 4-way repeated measures linear model with main effects frequency range (*range*; lo-vs. hi-range), physiological condition (*cond*; EO vs. EC), spatial filter (*filt*; CAR vs. SL) and cortical location (*ch*; FC2/FC1 vs. POz). Beyond assessing if spectral exponents differ in low- and high-frequency regimes (*range*), if closing the eyes had an effect on spectral slope (*cond*), if spatial filtering affects slope estimates (*filt*) and if spectral slopes differ over various cortical locations (*ch*), all (2-, 3- and 4-way) interaction levels were also considered to evaluate if these conditions affect each other. For all linear models, sphericity of the data was evaluated via Mauchly’s test, and in case of violation main and interaction effect *p*-values were corrected using the Greenhouse-Geisser method. In case of a significant main effect or *cond ×range* interaction, we performed post hoc pairwise comparisons using paired t-tests with Bonferroni correction. Pairwise comparisons were also performed for oscillatory alpha power between EO and EC states. In case of normally distributed data a paired t-test was used, while a Wilcoxon signed rank test otherwise. To assess if changes observed in spectral slopes are related to changes in resting alpha BLP. We characterized the plausible relationship between spectral slope (∆*β*_*lo*_ and ∆*β*_*hi*_) and resting alpha power 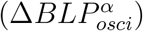 difference indices using Pearson or Spearman correlation, depending on data normality. In all analyses, outcomes were adjusted to multiple comparisons using the Bonferroni method at a significance level of *α* = 0.05. For pairwise comparisons, effect sizes were captured via Cohen’s *d*.

## Author Contributions

FSR: Conceptualization, Data curation, Formal analysis, Investigation, Methodology, Software, Visualization, Writing - original draft, Writing - review and editing; DMA: Data curation, Writing - review and editing; PM: Methodology, Software, Writing - review and editing; JM: Conceptualization, Investigation, Methodology, Writing - review and editing; JLC: Methodology, Writing - review and editing; JDRM: Conceptualization, Funding acquisition, Resources, Supervision, Writing - review and editing

## Code Availability, Data Availability and Online Supplementary Material

All custom scripts and functions, as well as supporting data will be made available for any interested researchers at the following GitHub repository: https://github.com/samuelracz/EEG_preproc_slope. This online repository also holds all main and supplementary figures generated throughout the analyses, as well as the codes to recreate them.

## References

[1] S. Ghatak, M. Talantova, S. R. McKercher, S. A. Lipton, Novel therapeutic approach for excitatory/inhibitory imbalance in neurodevelopmental and neurodegenerative diseases, Annual Review of Pharmacology and Toxicology 61 (1) (2021) 701–721.

[2] S. B. Nelson, V. Valakh, Excitatory/inhibitory balance and circuit homeostasis in autism spectrum disorders, Neuron 87 (4) (2015) 684–698.

[3] V. S. Sohal, J. L. Rubenstein, Excitation-inhibition balance as a framework for investigating mechanisms in neuropsychiatric disorders, Molecular psychiatry 24 (9) (2019) 1248–1257.

[4] Y. Liu, P. Ouyang, Y. Zheng, L. Mi, J. Zhao, Y. Ning, W. Guo, A selective review of the excitatory-inhibitory imbalance in schizophrenia: underlying biology, genetics, microcircuits, and symptoms, Frontiers in cell and developmental biology 9 (2021) 664535.

[5] F. Maestú, W. de Haan, M. A. Busche, J. DeFelipe, Neuronal excitation/inhibition imbalance: core element of a translational perspective on alzheimer pathophysiology, Ageing Research Reviews 69 (2021) 101372.

[6] S. Hafizi, T. K. Rajji, Modifiable risk factors of dementia linked to excitation-inhibition imbalance, Ageing Research Reviews 83 (2023) 101804.

[7] B. Haider, A. Duque, A. R. Hasenstaub, D. A. McCormick, Neocortical network activity in vivo is generated through a dynamic balance of excitation and inhibition, Journal of Neuroscience 26 (17) (2006) 4535–4545.

[8] R. Gao, E. J. Peterson, B. Voytek, Inferring synaptic excitation/inhibition balance from field potentials, Neuroimage 158 (2017) 70–78.

[9] B. J. He, Scale-free brain activity: past, present, and future, Trends in cognitive sciences 18 (9) (2014) 480–487.

[10] G. F. Grosu, A. V. Hopp, V. V. Moca, H. Bârzan, A. Ciuparu, M. Ercsey-Ravasz, M. Winkel, H. Linde, R. C. Mures, an, The fractal brain: scale-invariance in structure and dynamics, Cerebral Cortex 33 (8) (2023) 4574–4605.

[11] A. Eke, P. Herman, L. Kocsis, L. Kozak, Fractal characterization of complexity in temporal physiological signals, Physiological measurement 23 (1) (2002) R1.

[12] H. Wen, Z. Liu, Separating fractal and oscillatory components in the power spectrum of neurophysiological signal, Brain topography 29 (2016) 13–26.

[13] S. D. Muthukumaraswamy, D. T. Liley, 1/f electrophysiological spectra in resting and drug-induced states can be explained by the dynamics of multiple oscillatory relaxation processes, NeuroImage 179 (2018) 582–595.

[14] J. L. Molina, B. Voytek, M. L. Thomas, Y. B. Joshi, S. G. Bhakta, J. A. Talledo, N. R. Swerdlow, G. A. Light, Memantine effects on electroencephalographic measures of putative excitatory/inhibitory balance in schizophrenia, Biological Psychiatry: Cognitive Neuroscience and Neuroimaging 5 (6) (2020) 562–568.

[15] F. Akbarian, C. Rossi, L. Costers, M. B. D’hooghe, M. D’haeseleer, G. Nagels, J. Van Schependom, The spectral slope as a marker of excitation/inhibition ratio and cognitive functioning in multiple sclerosis, Human Brain Mapping 44 (17) (2023) 5784–5794.

[16] M. Chini, T. Pfeffer, I. Hanganu-Opatz, An increase of inhibition drives the developmental decorrelation of neural activity, Elife 11 (2022) e78811.

[17] P. Martínez-Cañada, E. Perez-Valero, J. Minguillon, F. Pelayo, M. A. López-Gordo, C. Morillas, Combining aperiodic 1/f slopes and brain simulation: An eeg/meg proxy marker of excitation/inhibition imbalance in alzheimer’s disease, Alzheimer’s & Dementia: Diagnosis, Assessment & Disease Monitoring 15 (3) (2023) e12477.

[18] T. Donoghue, M. Haller, E. J. Peterson, P. Varma, P. Sebastian, R. Gao, T. Noto, A. H. Lara, J. D. Wallis, R. T. Knight, et al., Parameterizing neural power spectra into periodic and aperiodic components, Nature neuroscience 23 (12) (2020) 1655–1665.

[19] V. Shirhatti, A. Borthakur, S. Ray, Effect of reference scheme on power and phase of the local field potential, Neural computation 28 (5) (2016) 882–913.

[20] R. Becker, D. Van De Ville, A. Kleinschmidt, Alpha oscillations reduce temporal long-range dependence in spontaneous human brain activity, Journal of Neuroscience 38 (3) (2018) 755–764.

[21] D. J. McFarland, L. M. McCane, S. V. David, J. R. Wolpaw, Spatial filter selection for eeg-based communication, Electroencephalography and clinical Neurophysiology 103 (3) (1997) 386–394.

[22] R. J. Barry, A. R. Clarke, S. J. Johnstone, C. A. Magee, J. A. Rushby, Eeg differences between eyes-closed and eyes-open resting conditions, Clinical neurophysiology 118 (12) (2007) 2765–2773.

[23] O. Jensen, A. Mazaheri, Shaping functional architecture by oscillatory alpha activity: gating by inhibition, Frontiers in human neuroscience 4 (2010) 186.

[24] K. J. Miller, L. B. Sorensen, J. G. Ojemann, M. Den Nijs, Power-law scaling in the brain surface electric potential, PLoS computational biology 5 (12) (2009) e1000609.

[25] B. J. He, J. M. Zempel, A. Z. Snyder, M. E. Raichle, The temporal structures and functional significance of scale-free brain activity, Neuron 66 (3) (2010) 353–369.

[26] F. S. Racz, K. Farkas, O. Stylianou, Z. Kaposzta, A. Czoch, P. Mukli, G. Csukly, A. Eke, Separating scale-free and oscillatory components of neural activity in schizophrenia, Brain and behavior 11 (5) (2021) e02047.

[27] Z. Nagy, P. Mukli, P. Herman, A. Eke, Decomposing multifractal crossovers, Frontiers in Physiology 8 (2017) 271083.

[28] B. Voytek, M. A. Kramer, J. Case, K. Q. Lepage, Z. R. Tempesta, R. T. Knight, A. Gazzaley, Age-related changes in 1/f neural electrophysiological noise, Journal of neuroscience 35 (38) (2015) 13257–13265.

[29] S. Dave, T. Brothers, T. Swaab, 1/f neural noise and electrophysiological indices of contextual prediction in aging, Brain research 1691 (2018) 34–43.

[30] A. Pathania, M. Euler, M. Clark, R. Cowan, K. Duff, K. Lohse, Resting eeg spectral slopes are associated with age-related differences in information processing speed, Biological Psychology 168 (2022) 108261.

[31] A. Czoch, Z. Kaposzta, P. Mukli, O. Stylianou, A. Eke, F. S. Racz, Resting-state fractal brain connectivity is associated with impaired cognitive performance in healthy aging, GeroScience 46 (1) (2024) 473–489.

[32] F. S. Racz, K. Farkas, M. Becske, H. Molnar, Z. Fodor, P. Mukli, G. Csukly, Reduced temporal variability of cortical excitation/inhibition ratio in schizophrenia, Schizophrenia 11 (1) (2025) 20.

[33] F. Lombardi, H. J. Herrmann, L. Parrino, D. Plenz, S. Scarpetta, A. E. Vaudano, L. De Arcangelis, O. Shriki, Beyond pulsed inhibition: Alpha oscillations modulate attenuation and amplification of neural activity in the awake resting state, Cell reports 42 (10) (2023).

[34] M. Sano, Y. Nishiura, I. Morikawa, A. Hoshino, J.-i. Uemura, K. Iwatsuki, H. Hirata, M. Hoshiyama, Analysis of the alpha activity envelope in electroencephalography in relation to the ratio of excitatory to inhibitory neural activity, PLoS One 19 (6) (2024) e0305082.

[35] D. Yao, L. Wang, R. Oostenveld, K. D. Nielsen, L. Arendt-Nielsen, A. C. Chen, A comparative study of different references for eeg spectral mapping: the issue of the neutral reference and the use of the infinity reference, Physiological measurement 26 (3) (2005) 173.

[36] X. Lei, K. Liao, Understanding the influences of eeg reference: a largescale brain network perspective, Frontiers in neuroscience 11 (2017) 205.

[37] N. Kozhemiako, D. Mylonas, J. Q. Pan, M. J. Prerau, S. Redline, S. M. Purcell, Sources of variation in the spectral slope of the sleep eeg, eneuro 9 (5) (2022).

[38] F. Perrin, O. Bertrand, J. Pernier, Scalp current density mapping: value and estimation from potential data, IEEE Transactions on biomedical engineering (4) (1987) 283–288.

[39] J. W. Kantelhardt, E. Koscielny-Bunde, H. H. Rego, S. Havlin, A. Bunde, Detecting long-range correlations with detrended fluctuation analysis, Physica A: Statistical Mechanics and its Applications 295 (3-4) (2001) 441–454.

[40] Z. Chen, P. C. Ivanov, K. Hu, H. E. Stanley, Effect of nonstationarities on detrended fluctuation analysis, Physical review E 65 (4) (2002) 041107.

[41] L. Kristoufek, Mixed-correlated arfima processes for power-law cross-correlations, Physica A: Statistical Mechanics and its Applications 392 (24) (2013) 6484–6493.

[42] J. Echeverria, E. Rodriguez, M. Aguilar-Cornejo, J. Alvarez-Ramirez, Linear combination of power-law functions for detecting multiscaling using detrended fluctuation analysis, Physica A: Statistical Mechanics and its Applications 460 (2016) 283–293.

[43] G. Chiarion, L. Sparacino, Y. Antonacci, L. Faes, L. Mesin, Connectivity analysis in eeg data: a tutorial review of the state of the art and emerging trends, Bioengineering 10 (3) (2023) 372.

[44] S. P. van den Broek, F. Reinders, M. Donderwinkel, M. Peters, Volume conduction effects in eeg and meg, Electroencephalography and clinical neurophysiology 106 (6) (1998) 522–534.

[45] Y. Yamamoto, R. L. Hughson, Coarse-graining spectral analysis: new method for studying heart rate variability, Journal of Applied Physiology 71 (3) (1991) 1143–1150.

[46] A. Eke, P. Hermán, M. Hajnal, Fractal and noisy cbv dynamics in humans: influence of age and gender, Journal of Cerebral Blood Flow & Metabolism 26 (7) (2006) 891–898.

[47] P. Herman, B. G. Sanganahalli, F. Hyder, A. Eke, Fractal analysis of spontaneous fluctuations of the bold signal in rat brain, Neuroimage 58 (4) (2011) 1060–1069.

[48] G. Pellegrino, T. Hedrich, V. Sziklas, J.-M. Lina, C. Grova, E. Kobayashi, How cerebral cortex protects itself from interictal spikes: The alpha/beta inhibition mechanism, Human brain mapping 42 (11) (2021) 3352–3365.

[49] B. Krause, J. Márquez-Ruiz, R. C. Kadosh, The effect of transcranial direct current stimulation: a role for cortical excitation/inhibition balance?, Frontiers in human neuroscience 7 (2013) 602.

[50] O. Lányi, B. Koleszár, A. Schulze Wenning, D. Balogh, M. A. Engh, A. A. Horváth, P. Fehérvari, P. Hegyi, Z. Molnár, Z. Unoka, et al., Excitation/inhibition imbalance in schizophrenia: a meta-analysis of inhibitory and excitatory tms-emg paradigms, Schizophrenia 10 (1) (2024) 56.

[51] H. Alawieh, D. Liu, J. Madera, S. Kumar, F. S. Racz, A. M. Fey, J. d. R. Millán, Transcutaneous electrical spinal cord stimulation promotes focal sensorimotor activation that accelerates brain-computer interface skill learning, medRxiv (2024) 2024–06.

[52] O. Bazanova, D. Vernon, Interpreting eeg alpha activity, Neuroscience & Biobehavioral Reviews 44 (2014) 94–110.

[53] Y. Takei, K. Fujihara, M. Tagawa, N. Hironaga, J. Near, M. Kasagi, Y. Takahashi, T. Motegi, Y. Suzuki, Y. Aoyama, et al., The inhibition/excitation ratio related to task-induced oscillatory modulations during a working memory task: a multtimodal-imaging study using meg and mrs, Neuroimage 128 (2016) 302–315.

[54] C. Carvalhaes, J. A. De Barros, The surface laplacian technique in eeg: Theory and methods, International Journal of Psychophysiology 97 (3) (2015) 174–188.

[55] B. B. Mandelbrot, J. W. Van Ness, Fractional brownian motions, fractional noises and applications, SIAM review 10 (4) (1968) 422–437.

[56] Y. Yamamoto, R. L. Hughson, Extracting fractal components from time series, Physica D: Nonlinear Phenomena 68 (2) (1993) 250–264.

[57] F. S. Racz, A. Czoch, Z. Kaposzta, O. Stylianou, P. Mukli, A. Eke, Multiple-resampling cross-spectral analysis: an unbiased tool for estimating fractal connectivity with an application to neurophysiological signals, Frontiers in Physiology 13 (2022) 817239.

