## Supplementary Material for "Alternative EEG pre-processing pipelines can lead to conflicting conclusions regarding cortical excitation/inhibition ratio"

[a] *Department of Neurology, The University of Texas at Austin, 1601 Trinity Street, Austin, 78712, TX, United States*
[b] *Mulva Clinic for the Neurosciences, The University of Texas at Austin, 1601 Trinity Street, Austin, 78712, TX, United States*
[c] *Chandra Family Department of Electrical and Computer Engineering, The University of Texas at Austin, 2501 Speedway, Austin, 78712, TX, United States*
[d] *Oklahoma Center for Geroscience and Healthy Brain Aging, University of Oklahoma Health Sciences Center, 1100 N Lindsay Ave, Oklahoma City, 73104, OK, United States*
[e] *Department of Neurosurgery, University of Oklahoma Health Sciences Center, 1000 N Lincoln Blvd, Oklahoma City, 73104, OK, United States*
[f] *Centro de Física, Instituto Venezolano de Investigaciones Científicas, 92WF+CG4, Caracas, 1020-A, Miranda, Venezuela*
[g] *Departamento de Física Aplicada, ETSI Aeronáutica y del Espacio, Universidad Politécnica de Madrid, Pza. Cardenal Cisneros 3, Madrid, 28040, , Spain*

### Supplementary Figures and Tables

*Corresponding author: Frigyes Samuel Racz

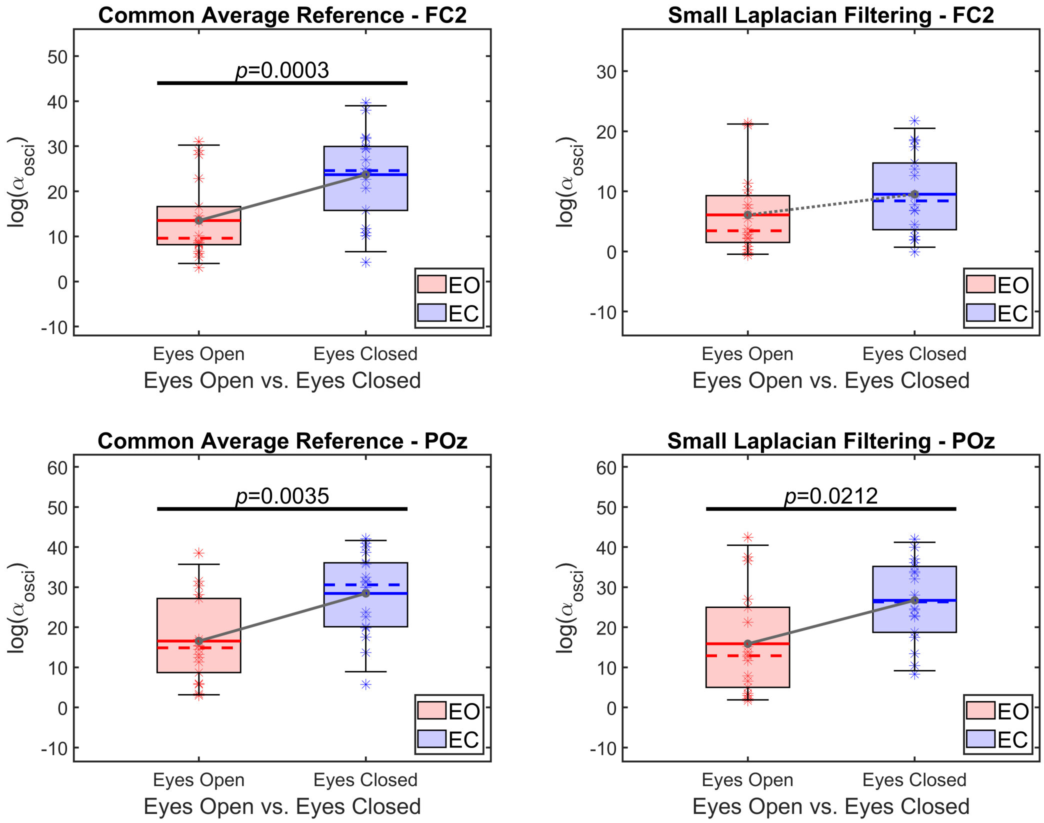


Figure 1: Isolated alpha band limited power in eyes open (blue) vs. eyes closed (red) states over FC2 (upper panels) and POz (lower panels). Left and right columns show CAR- and SL-filtered data, respectively. Horizontal black line indicates significant between-state difference. EO: eyes open; EC: eyes closed; CAR: common average reference; SL: small Laplacian filtering.

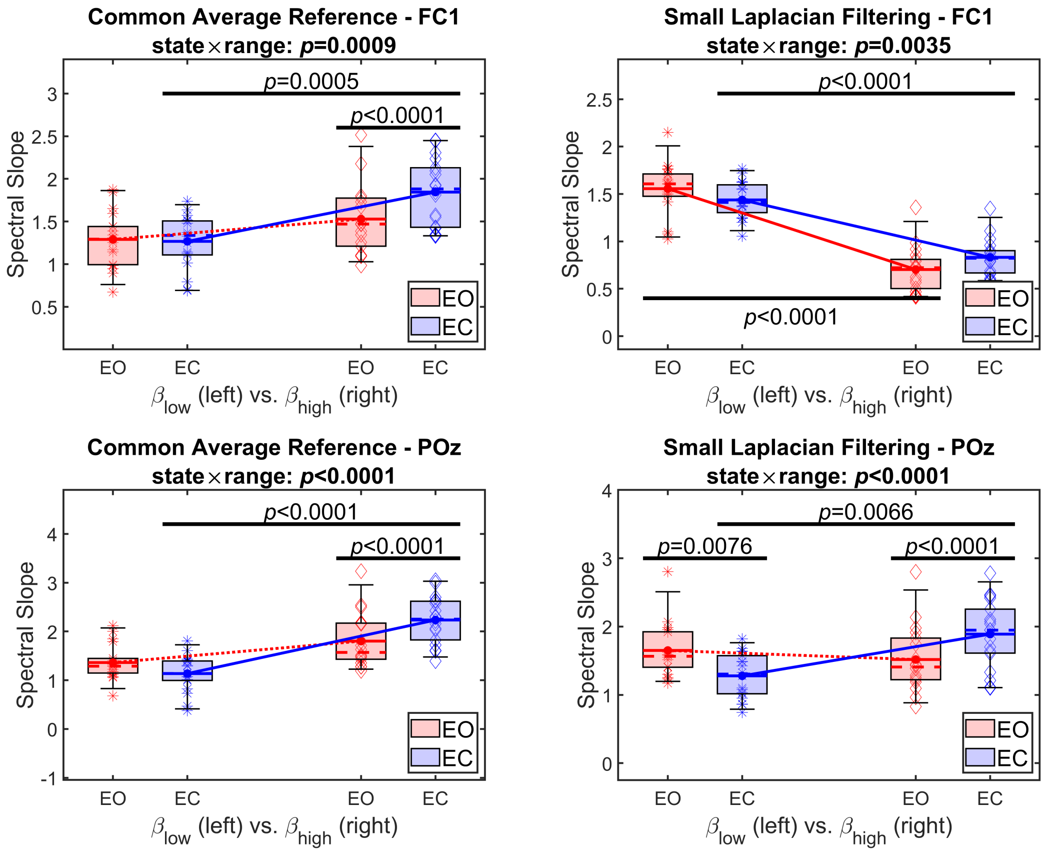


Figure 2: Summary of results from the spectral slope analysis in Cohort #1 including channels FC1 and POz for laterality control. Left and right panels show outcomes from CAR and SL pipelines, while upper and lower rows depict results from FC1 and POz, respectively. Box plots for $\beta$ estimates from EO are denoted in red, while those from EC in blue. On every panel, box plots on the left illustrate $\beta_{lo}$, while those on the right $\beta_{hi}$. Significant pairwise differences are denoted by black vertical bars, and the $p$-value for the $state \times range$ interaction effect is indicated in the panel title. CAR: Common Average Reference; SL: Small Laplacian; EO: Eyes Open; EC: Eyes Closed.

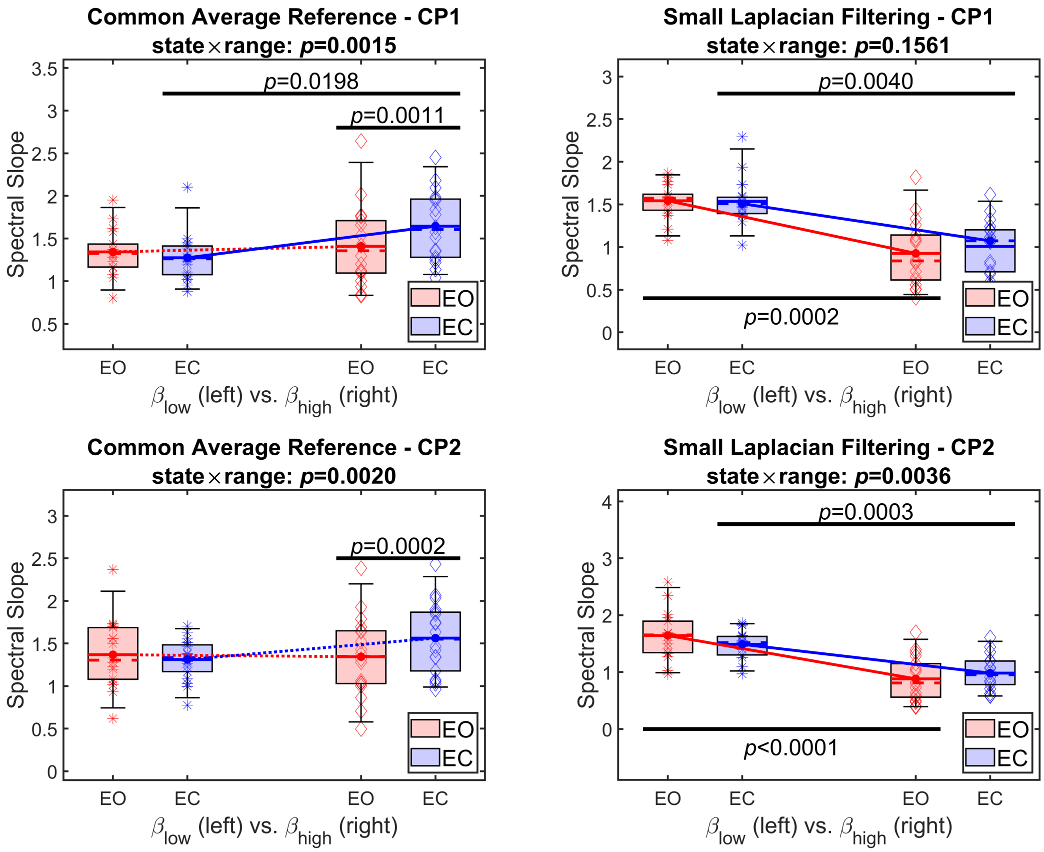


Figure 3: Summary of results from the spectral slope analysis in Cohort #1 including channels CP1 and CP2. Left and right panels show outcomes from CAR and SL pipelines, while upper and lower rows depict results from CP1 and CP2, respectively. Box plots for $\beta$ estimates from EO are denoted in red, while those from EC in blue. On every panel, box plots on the left illustrate $\beta_{lo}$, while those on the right $\beta_{hi}$. Significant pairwise differences are denoted by black vertical bars, and the $p$-value for the $state \times range$ interaction effect is indicated in the panel title. CAR: Common Average Reference; SL: Small Laplacian; EO: Eyes Open; EC: Eyes Closed.

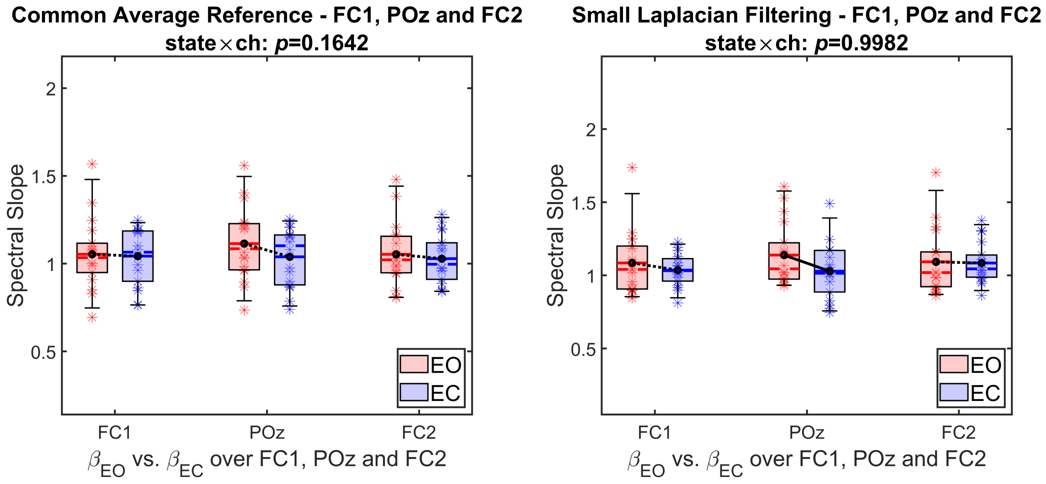


Figure 4: Summary of results from broadband ($1-45$ Hz) spectral slope analysis in Cohort #1 including channels FC1, POz and FC2. Left and right panels show outcomes from CAR and SL pipelines, respectively. Box plots for $\beta$ estimates from EO are denoted in red, while those from EC in blue. On every panel, box plots from left to right illustrate $\beta_{bb}$ over FC1, POz and FC2. Significant pairwise differences are denoted by black vertical bars, and the $p$-value for the *state* $\times$ *range* interaction effect is indicated in the panel title. CAR: Common Average Reference; SL: Small Laplacian; EO: Eyes Open; EC: Eyes Closed.

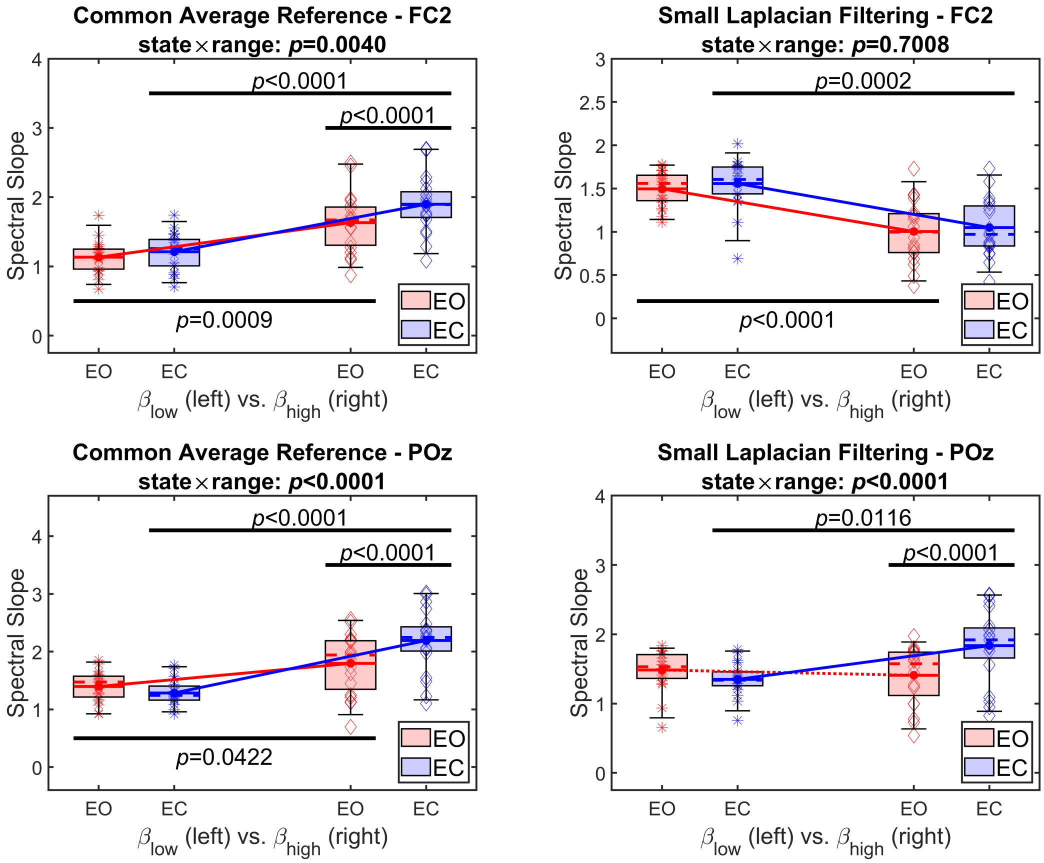


Figure 5: Summary of results from the spectral slope analysis in Cohort #2 including channels FC2 and POz. Left and right panels show outcomes from CAR and SL pipelines, while upper and lower rows depict results from FC2 and POz, respectively. Box plots for $\beta$ estimates from EO are denoted in red, while those from EC in blue. On every panel, box plots on the left illustrate $\beta_{lo}$, while those on the right $\beta_{hi}$. Significant pairwise differences are denoted by black vertical bars, and the $p$-value for the $state \times range$ interaction effect is indicated in the panel title. CAR: Common Average Reference; SL: Small Laplacian; EO: Eyes Open; EC: Eyes Closed.

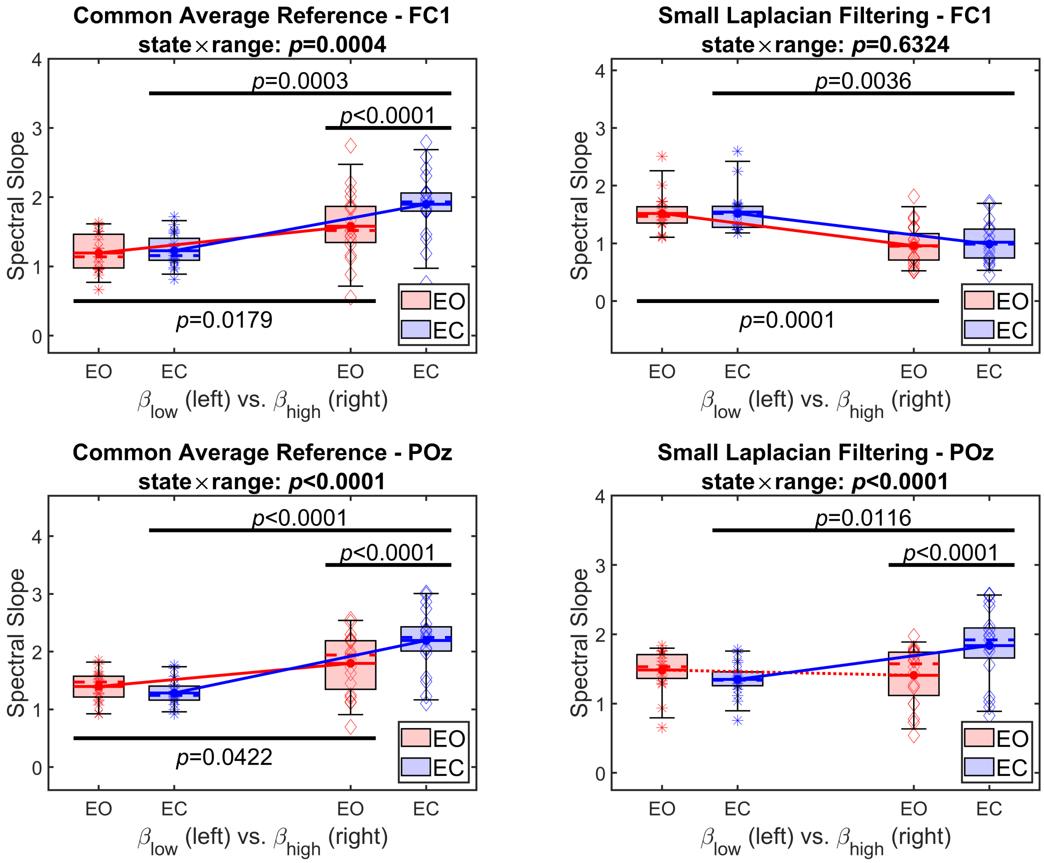


Figure 6: Summary of results from the spectral slope analysis in Cohort #2 including channels FC1 and POz for laterality control. Left and right panels show outcomes from CAR and SL pipelines, while upper and lower rows depict results from FC1 and POz, respectively. Box plots for $\beta$ estimates from EO are denoted in red, while those from EC in blue. On every panel, box plots on the left illustrate $\beta_{lo}$, while those on the right $\beta_{hi}$. Significant pairwise differences are denoted by black vertical bars, and the $p$-value for the $state \times range$ interaction effect is indicated in the panel title. CAR: Common Average Reference; SL: Small Laplacian; EO: Eyes Open; EC: Eyes Closed.

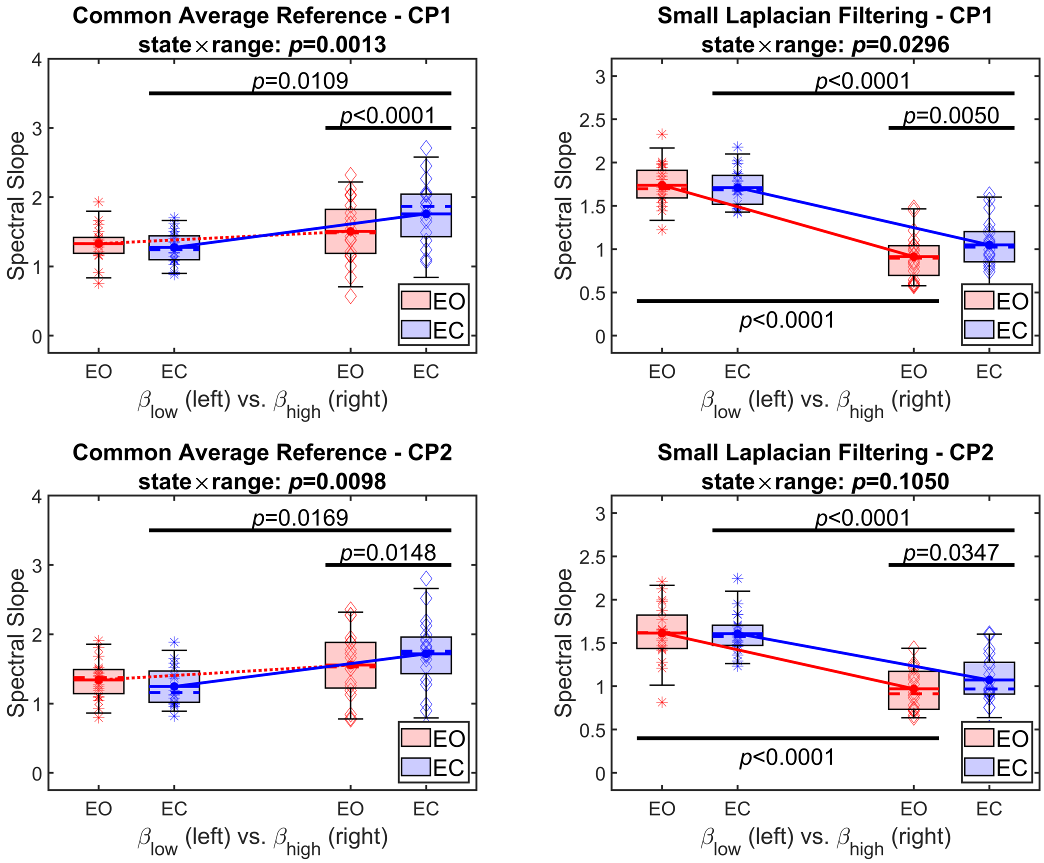


Figure 7: Summary of results from the spectral slope analysis in Cohort #2 including channels CP1 and CP2. Left and right panels show outcomes from CAR and SL pipelines, while upper and lower rows depict results from CP1 and CP2, respectively. Box plots for $\beta$ estimates from EO are denoted in red, while those from EC in blue. On every panel, box plots on the left illustrate $\beta_{lo}$, while those on the right $\beta_{hi}$. Significant pairwise differences are denoted by black vertical bars, and the $p$-value for the $state \times range$ interaction effect is indicated in the panel title. CAR: Common Average Reference; SL: Small Laplacian; EO: Eyes Open; EC: Eyes Closed.

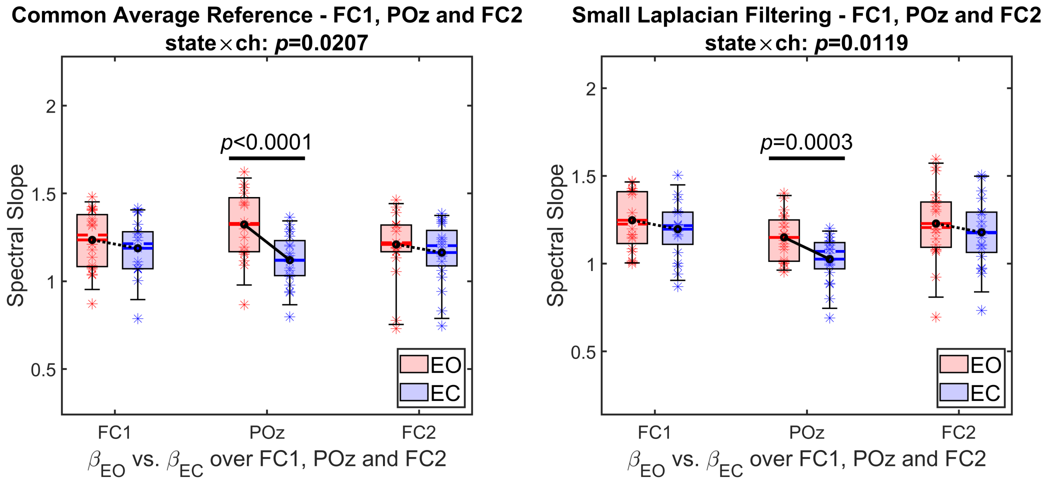


Figure 8: Summary of results from broadband (1 − 45 Hz) spectral slope analysis in Cohort #2 including channels FC1, POz and FC2. Left and right panels show outcomes from CAR and SL pipelines, respectively. Box plots for $\beta$ estimates from EO are denoted in red, while those from EC in blue. On every panel, box plots from left to right illustrate $\beta_{bb}$ over FC1, POz and FC2. Significant pairwise differences are denoted by black vertical bars, and the $p$-value for the *state* $\times$ *range* interaction effect is indicated in the panel title. CAR: Common Average Reference; SL: Small Laplacian; EO: Eyes Open; EC: Eyes Closed.

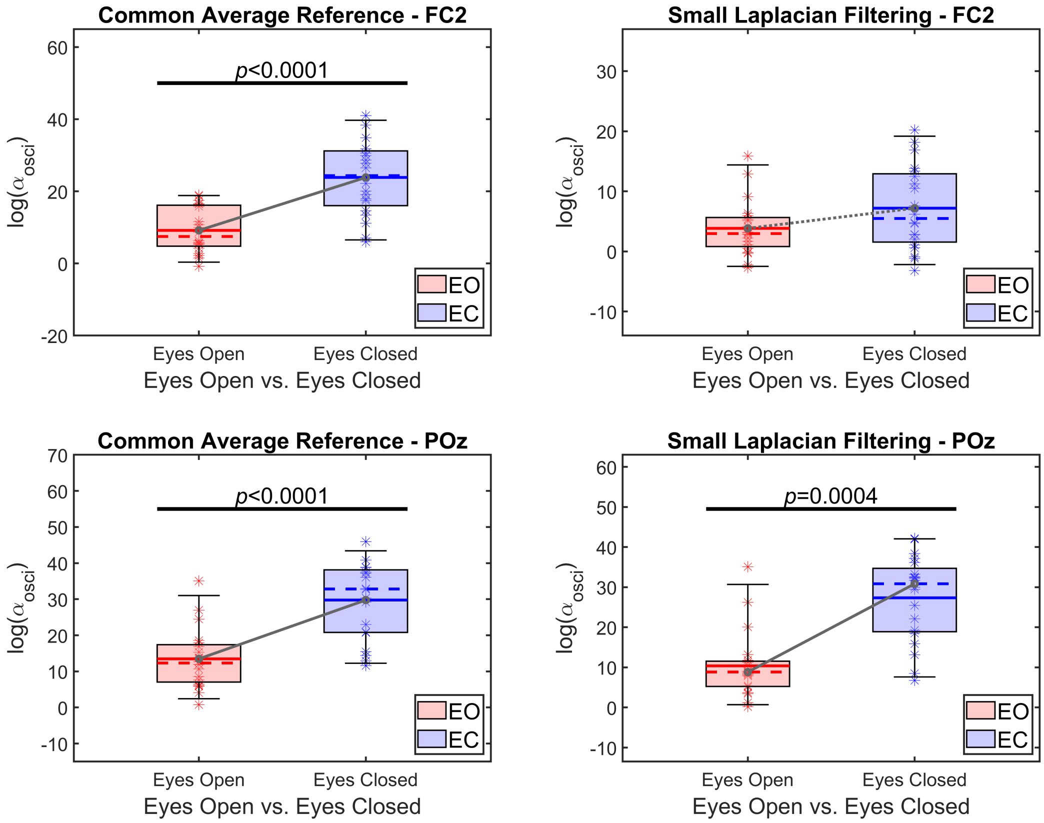


Figure 9: Isolated alpha band limited power in eyes open (blue) vs. eyes closed (red) states over FC2 (upper panels) and POz (lower panels) in Cohort #2. Left and right columns show CAR- and SL-filtered data, respectively. Horizontal black line indicates significant between-state difference. EO: eyes open; EC: eyes closed; CAR: common average reference; SL: small Laplacian filtering.

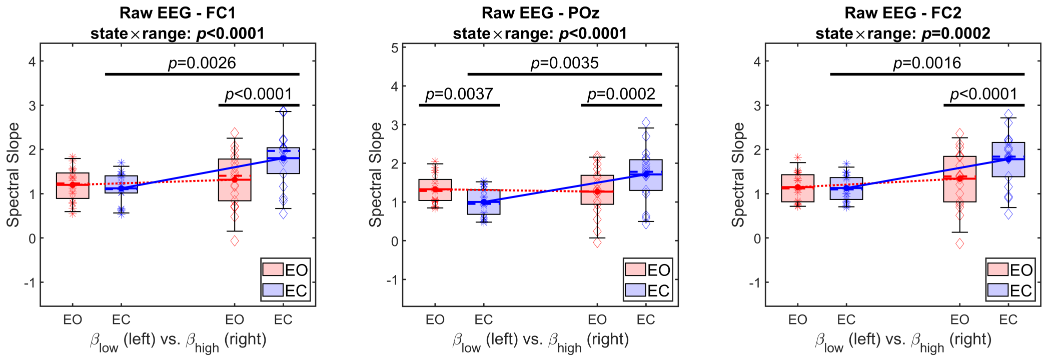


Figure 10: Summary of results from the spectral slope analysis of native/raw EEG data in Cohort #1 including channels FC1, POz and FC2. Panels from left to right show outcomes for FC1, POz and FC2. Box plots for $\beta$ estimates from EO are denoted in red, while those from EC in blue. On every panel, box plots on the left illustrate $\beta_{lo}$, while those on the right $\beta_{hi}$. Significant pairwise differences are denoted by black vertical bars, and the $p$-value for the $state \times range$ interaction effect is indicated in the panel title. CAR: Common Average Reference; SL: Small Laplacian; EO: Eyes Open; EC: Eyes Closed.

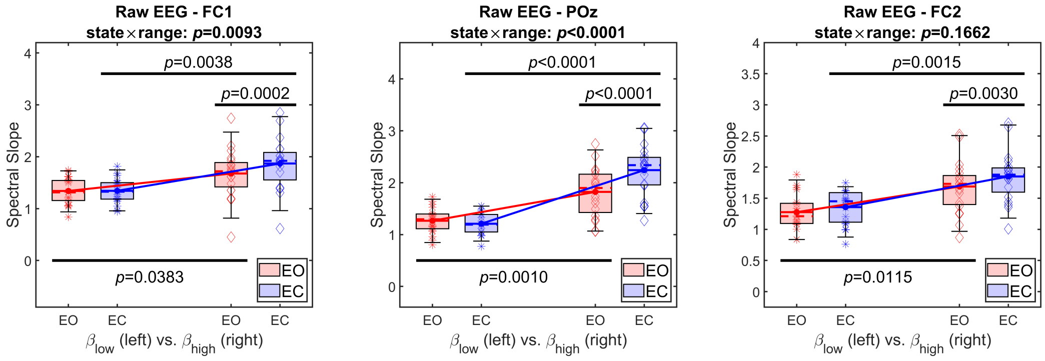


Figure 11: Summary of results from the spectral slope analysis of native/raw EEG data in Cohort #2 including channels FC1, POz and FC2. Panels from left to right show outcomes for FC1, POz and FC2. Box plots for $\beta$ estimates from EO are denoted in red, while those from EC in blue. On every panel, box plots on the left illustrate $\beta_{lo}$, while those on the right $\beta_{hi}$. Significant pairwise differences are denoted by black vertical bars, and the $p$-value for the $state \times range$ interaction effect is indicated in the panel title. CAR: Common Average Reference; SL: Small Laplacian; EO: Eyes Open; EC: Eyes Closed.

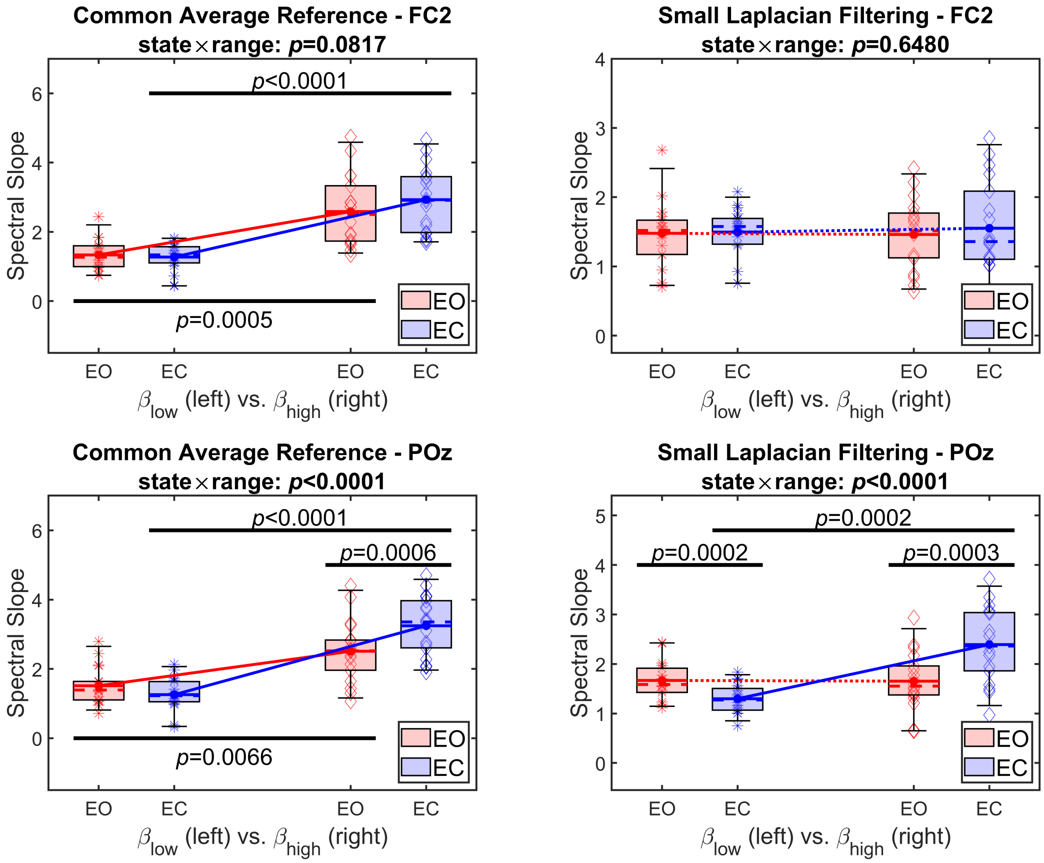


Figure 12: Summary of results from FOOOF in Cohort #1 including channels FC2 and POz. Left and right panels show outcomes from CAR and SL pipelines, while upper and lower rows depict results from FC2 and POz, respectively. Box plots for $\beta$ estimates from EO are denoted in red, while those from EC in blue. On every panel, box plots on the left illustrate $\beta_{lo}$, while those on the right $\beta_{hi}$. Significant pairwise differences are denoted by black vertical bars, and the $p$-value for the $state \times range$ interaction effect is indicated in the panel title. FOOOF: fitting oscillations and one over f; CAR: Common Average Reference; SL: Small Laplacian; EO: Eyes Open; EC: Eyes Closed.

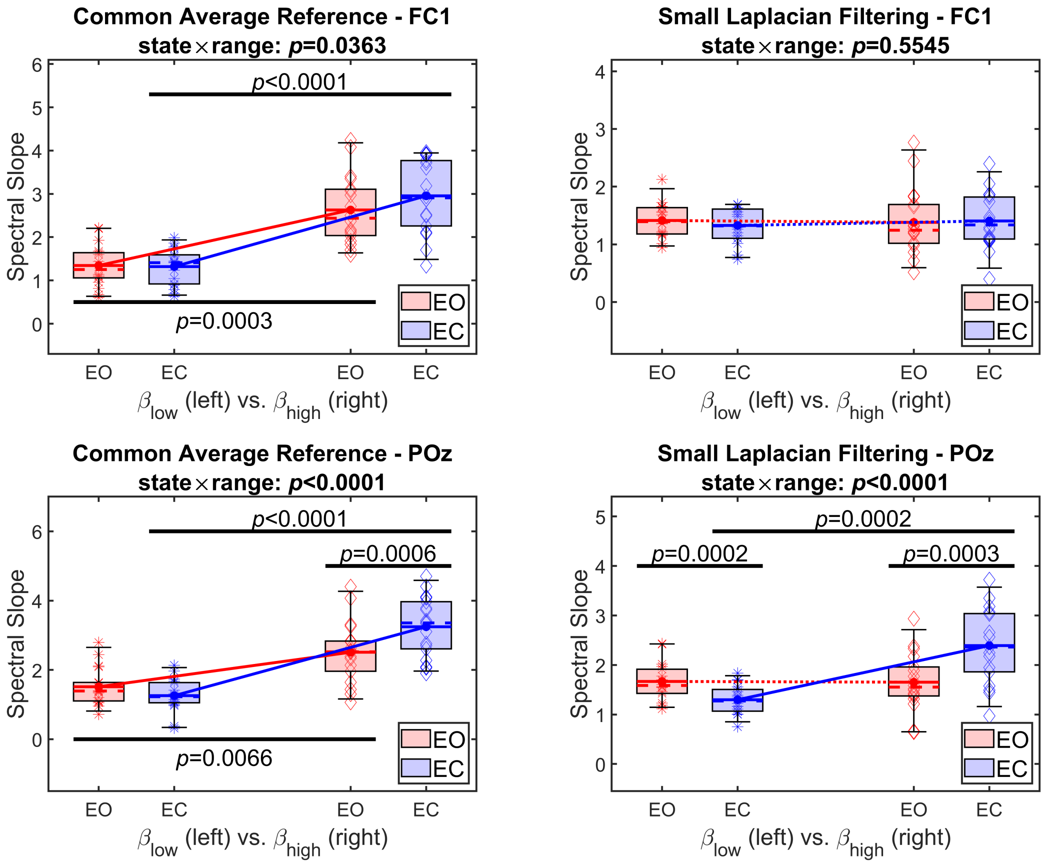


Figure 13: Summary of results from FOOOF in Cohort #1 including channels FC1 and POz for laterality control. Left and right panels show outcomes from CAR and SL pipelines, while upper and lower rows depict results from FC1 and POz, respectively. Box plots for $\beta$ estimates from EO are denoted in red, while those from EC in blue. On every panel, box plots on the left illustrate $\beta_{lo}$, while those on the right $\beta_{hi}$. Significant pairwise differences are denoted by black vertical bars, and the $p$-value for the $state \times range$ interaction effect is indicated in the panel title. FOOOF: fitting oscillations and one over f; CAR: Common Average Reference; SL: Small Laplacian; EO: Eyes Open; EC: Eyes Closed.

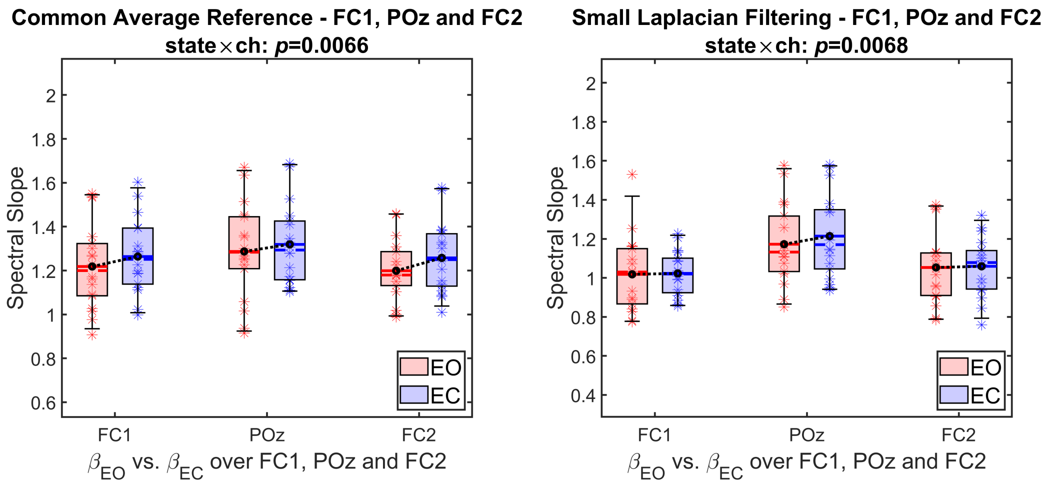


Figure 14: Summary of results from broadband ($1 - 45$ Hz) FOOOF analysis in Cohort #1 including channels FC1, POz and FC2. Left and right panels show outcomes from CAR and SL pipelines, respectively. Box plots for $\beta$ estimates from EO are denoted in red, while those from EC in blue. On every panel, box plots from left to right illustrate $\beta_{bb}$ over FC1, POz and FC2. Significant pairwise differences are denoted by black vertical bars, and the $p$-value for the $state \times range$ interaction effect is indicated in the panel title. FOOOF: fitting oscillations and one over f'; CAR: Common Average Reference; SL: Small Laplacian; EO: Eyes Open; EC: Eyes Closed.

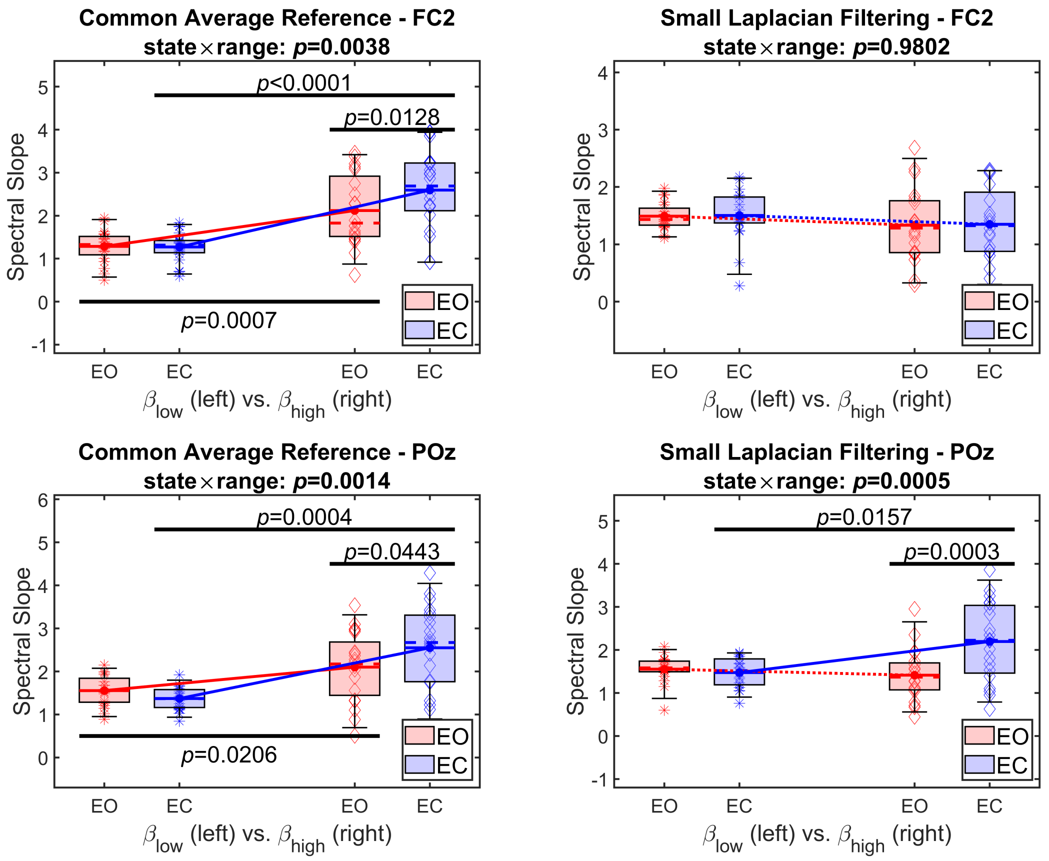


Figure 15: Summary of results from FOOOF in Cohort #2 including channels FC2 and POz. Left and right panels show outcomes from CAR and SL pipelines, while upper and lower rows depict results from FC2 and POz, respectively. Box plots for $\beta$ estimates from EO are denoted in red, while those from EC in blue. On every panel, box plots on the left illustrate $\beta_{lo}$, while those on the right $\beta_{hi}$. Significant pairwise differences are denoted by black vertical bars, and the $p$-value for the *state* $\times$ *range* interaction effect is indicated in the panel title. FOOOF: fitting oscillations and one over f; CAR: Common Average Reference; SL: Small Laplacian; EO: Eyes Open; EC: Eyes Closed.

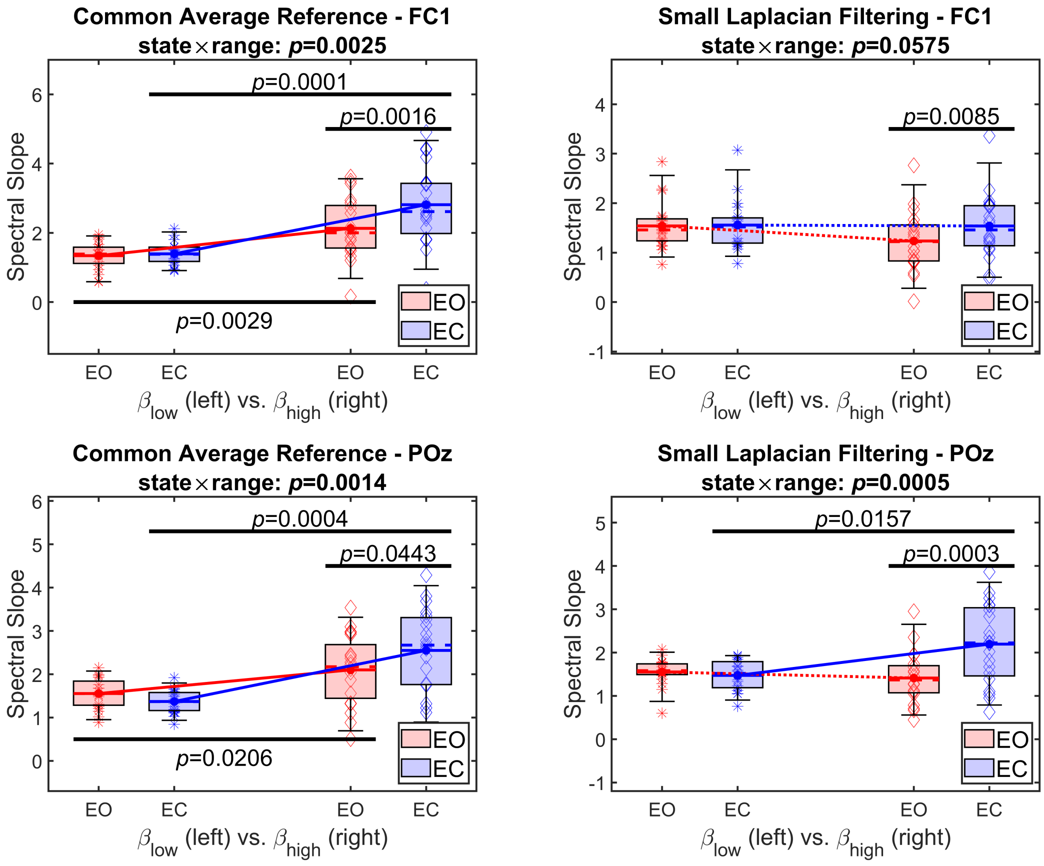


Figure 16: Summary of results from FOOOF in Cohort #2 including channels FC1 and POz for laterality control. Left and right panels show outcomes from CAR and SL pipelines, while upper and lower rows depict results from FC1 and POz, respectively. Box plots for $\beta$ estimates from EO are denoted in red, while those from EC in blue. On every panel, box plots on the left illustrate $\beta_{lo}$, while those on the right $\beta_{hi}$. Significant pairwise differences are denoted by black vertical bars, and the $p$-value for the $state \times range$ interaction effect is indicated in the panel title. FOOOF: fitting oscillations and one over f; CAR: Common Average Reference; SL: Small Laplacian; EO: Eyes Open; EC: Eyes Closed.

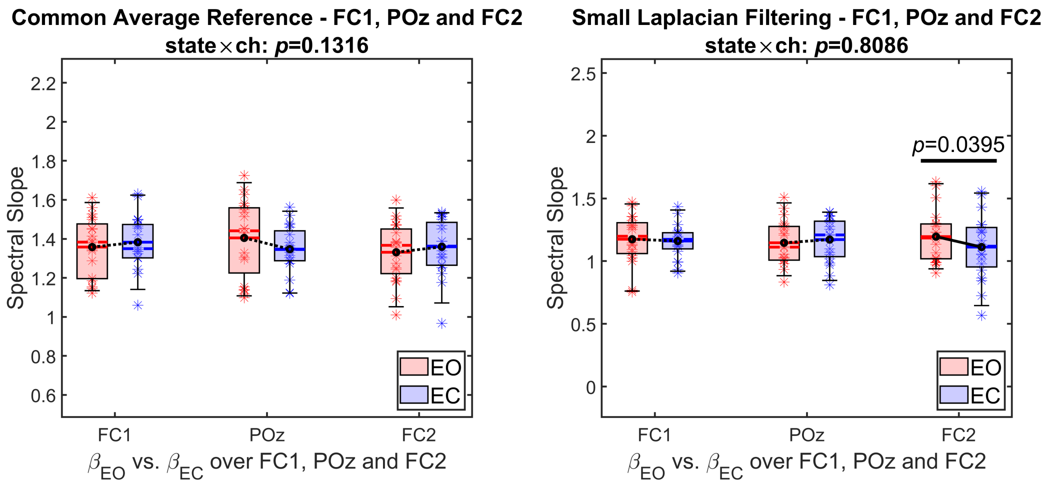


Figure 17: Summary of results from broadband ($1-45$ Hz) FOOOF analysis in Cohort #2 including channels FC1, POz and FC2. Left and right panels show outcomes from CAR and SL pipelines, respectively. Box plots for $\beta$ estimates from EO are denoted in red, while those from EC in blue. On every panel, box plots from left to right illustrate $\beta_{bb}$ over FC1, POz and FC2. Significant pairwise differences are denoted by black vertical bars, and the $p$-value for the $state \times range$ interaction effect is indicated in the panel title. FOOOF: fitting oscillations and one over f'; CAR: Common Average Reference; SL: Small Laplacian; EO: Eyes Open; EC: Eyes Closed.

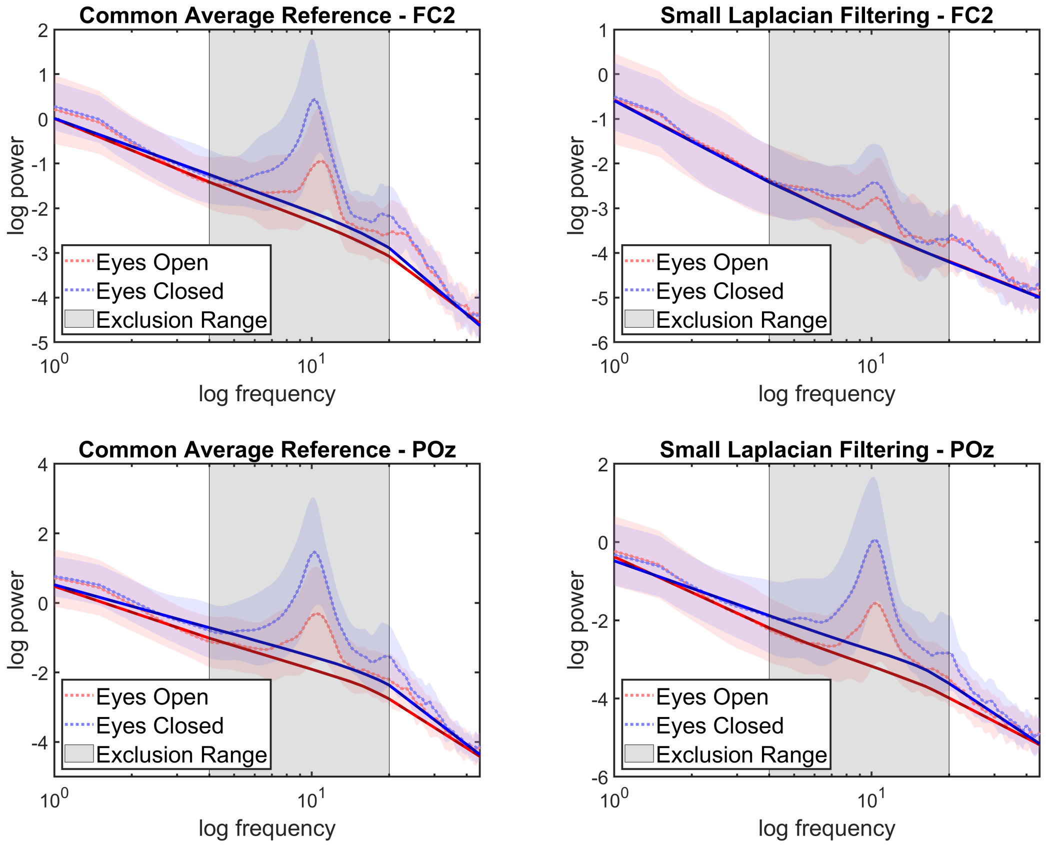


Figure 18: Illustration of the MMSPM method via group-averaged power spectra obtained from Cohort #1, using CAR- (left panels) and SL-treated (right panels) EEG data over FC2 (upper row) and POz (lower row). Dotted and continuous lines denote the raw/native and isolated $1/f$ spectra, respectively, while shaded, colored areas mark the corresponding standard error of the mean. The gray area indicates the frequency range excluded from slope estimation ($4-20$ Hz). The piece-wise linear model was fitted assuming two scaling ranges with an unknown breakpoint between 4-20 Hz. The inversion of $\beta_{lo} < \beta_{hi}$ pattern is apparent in the upper right panel (Fc2, SL-treated data). CAR: Common Average Reference; SL: Small Laplacian; MMSPM: MultiModal Spectral Parametrization Method.

Table 1: 4-way repeated measures ANOVA in Cohort #1. Included channels are FC2 and POz. Bold indicates significant main effect or interaction. *d*: degrees of freedom; SumSq: sum of squared errors; GG: Greenhouse-Geisser correction; *filt*: spatial filter; *ch*: EEG channel; *cond*: physiological condition; *range*: frequency range for slope estimation.

| Effect | $d$ | MeanSq | F | $p$-value (GG) |
|---|---|---|---|---|
| $filt$ | 17 | 1.9179 | 46.114 | $\mathbf{3.1488 \times 10^{-6}}$ |
| $ch$ | 17 | 4.7635 | 27.048 | $\mathbf{7.2156 \times 10^{-5}}$ |
| $cond$ | 17 | 0.28562 | 1.5617 | 0.22835 |
| $range$ | 17 | 3.3884 | 3.826 | 0.067102 |
| $filt \times ch$ | 17 | 0.9246 | 27.561 | $\mathbf{6.5194 \times 10^{-5}}$ |
| $filt \times cond$ | 17 | 0.17435 | 5.9852 | **0.025597** |
| $ch \times cond$ | 17 | 0.009119 | 0.36637 | 0.55298 |
| $filt \times range$ | 17 | 11.038 | 141.54 | $\mathbf{1.1478 \times 10^{-9}}$ |
| $ch \times range$ | 17 | 5.9352 | 26.877 | $\mathbf{7.466 \times 10^{-5}}$ |
| $cond \times range$ | 17 | 3.4358 | 38.07 | $\mathbf{1.0275 \times 10^{-5}}$ |
| $filt \times ch \times cond$ | 17 | 0.00043 | 0.0236 | 0.87977 |
| $filt \times ch \times range$ | 17 | 1.1514 | 31.526 | $\mathbf{3.093 \times 10^{-5}}$ |
| $filt \times cond \times range$ | 17 | 0.0741 | 4.0745 | 0.059597 |
| $ch \times cond \times range$ | 17 | 1.2527 | 36.435 | $\mathbf{1.3356 \times 10^{-5}}$ |
| $filt \times ch \times cond \times range$ | 17 | 0.2067 | 9.7464 | **0.006206** |

Table 2: 3-way repeated measures ANOVA in Cohort #1. for Common Average Reference filtering. Included channels are FC2 and POz. Bold indicates significant main effect or interaction. *d*: degrees of freedom; SumSq: sum of squared errors; GG: Greenhouse-Geisser correction; *ch*: EEG channel; *cond*: physiological condition; *range*: frequency range for slope estimation.

| Effect | $d$ | SumSq | F | $p$-value (GG) |
|---|---|---|---|---|
| $ch$ | 17 | 0.74541 | 12.209 | **0.00278** |
| $cond$ | 17 | 0.45315 | 2.9803 | 0.10241 |
| $range$ | 17 | 13.329 | 21.905 | **0.000215** |
| $ch \times cond$ | 17 | 0.002798 | 0.2048 | 0.65659 |
| $ch \times range$ | 17 | 0.92918 | 14.608 | **0.0013641** |
| $cond \times range$ | 17 | 2.2593 | 32.96 | $\mathbf{2.3989 \times 10^{-6}}$ |
| $ch \times cond \times range$ | 17 | 0.22083 | 19.857 | **0.00035** |

Table 3: 3-way repeated measures ANOVA in Cohort #1. for Small Laplacian filtering. Included channels are FC2 and POz. Bold indicates significant main effect or interaction. $d$: degrees of freedom; SumSq: sum of squared errors; GG: Greenhouse-Geisser correction; $ch$: EEG channel; $cond$: physiological condition; $range$: frequency range for slope estimation.

| Effect | $d$ | SumSq | F | $p$-value (GG) |
|---|---|---|---|---|
| $ch$ | 17 | 4.9427 | 33.26 | $\mathbf{2.2769 \times 10^{-5}}$ |
| $cond$ | 17 | 0.00683 | 0.11389 | 0.73989 |
| $range$ | 17 | 1.0975 | 3.0904 | 0.096747 |
| $ch \times cond$ | 17 | 0.00675 | 0.22967 | 0.63788 |
| $ch \times range$ | 17 | 6.1574 | 31.781 | $\mathbf{2.9547 \times 10^{-5}}$ |
| $cond \times range$ | 17 | 1.2505 | 31.36 | $\mathbf{3.1869 \times 10^{-6}}$ |
| $ch \times cond \times range$ | 17 | 1.2385 | 27.852 | $\mathbf{6.1569 \times 10^{-5}}$ |

Table 4: 4-way repeated measures ANOVA in Cohort #2. Included channels are FC2 and POz. Bold indicates significant main effect or interaction. $d$: degrees of freedom; SumSq: sum of squared errors; GG: Greenhouse-Geisser correction; $filt$: spatial filter; $ch$: EEG channel; $cond$: physiological condition; $range$: frequency range for slope estimation.

| Effect | $d$ | MeanSq | F | $p$-value (GG) |
|---|---|---|---|---|
| $filt$ | 19 | 2.3126 | 44.528 | $\mathbf{2.216 \times 10^{-6}}$ |
| $ch$ | 19 | 3.8796 | 23.626 | **0.0001** |
| $cond$ | 19 | 1.3212 | 31.5 | $\mathbf{2.063 \times 10^{-5}}$ |
| $range$ | 19 | 4.5197 | 5.4301 | 0.030967 |
| $filt \times ch$ | 19 | 0.040811 | 1.1334 | 0.30039 |
| $filt \times cond$ | 19 | 0.064644 | 5.4471 | **0.030736** |
| $ch \times cond$ | 19 | 0.02003 | 0.76425 | 0.39292 |
| $filt \times range$ | 19 | 11.89 | 167.08 | $\mathbf{7.3118 \times 10^{-11}}$ |
| $ch \times range$ | 19 | 2.9502 | 22.355 | **0.00015** |
| $cond \times range$ | 19 | 1.9317 | 39.307 | $\mathbf{5.0904 \times 10^{-6}}$ |
| $filt \times ch \times cond$ | 19 | 0.075189 | 10.99 | **0.00364** |
| $filt \times ch \times range$ | 19 | 2.0715 | 28.914 | $\mathbf{3.4476 \times 10^{-5}}$ |
| $filt \times cond \times range$ | 19 | 0.02766 | 2.0746 | 0.16605 |
| $ch \times cond \times range$ | 19 | 1.0158 | 33.038 | $\mathbf{1.5396 \times 10^{-5}}$ |
| $filt \times ch \times cond \times range$ | 19 | 0.083575 | 7.9747 | **0.010842** |

Table 5: 3-way repeated measures ANOVA in Cohort #2. for Common Average Reference filtering. Included channels are FC2 and POz. Bold indicates significant main effect or interaction. *d*: degrees of freedom; SumSq: sum of squared errors; GG: Greenhouse-Geisser correction; *ch*: EEG channel; *cond*: physiological condition; *range*: frequency range for slope estimation.

| Effect | $d$ | SumSq | F | $p$-value (GG) |
|---|---|---|---|---|
| $ch$ | 19 | 1.5623 | 23.823 | **0.0001** |
| $cond$ | 19 | 0.98515 | 25.397 | $\mathbf{7.2786 \times 10^{-5}}$ |
| $range$ | 19 | 15.536 | 28.191 | $\mathbf{4.0013 \times 10^{-5}}$ |
| $ch \times cond$ | 19 | 0.0088019 | 0.70879 | 0.41032 |
| $ch \times range$ | 19 | 0.038739 | 0.50463 | 0.4861 |
| $cond \times range$ | 19 | 1.2108 | 34.706 | $\mathbf{1.1322 \times 10^{-5}}$ |
| $ch \times cond \times range$ | 19 | 0.25833 | 17.41 | **0.00052** |

Table 6: 3-way repeated measures ANOVA in Cohort #2. for Small Laplacian filtering. Included channels are FC2 and POz. Bold indicates significant main effect or interaction. *d*: degrees of freedom; SumSq: sum of squared errors; GG: Greenhouse-Geisser correction; *ch*: EEG channel; *cond*: physiological condition; *range*: frequency range for slope estimation.

| Effect | $d$ | SumSq | F | $p$-value (GG) |
|---|---|---|---|---|
| $ch$ | 19 | 2.3581 | 17.515 | 0.0005 |
| $cond$ | 19 | 0.40066 | 26.676 | $\mathbf{5.5087 \times 10^{-5}}$ |
| $range$ | 19 | 0.87421 | 2.4807 | 0.13176 |
| $ch \times cond$ | 19 | 0.086418 | 4.1884 | 0.054791 |
| $ch \times range$ | 19 | 4.983 | 39.283 | $\mathbf{5.1112 \times 10^{-6}}$ |
| $cond \times range$ | 19 | 0.74853 | 27.134 | $\mathbf{4.9954 \times 10^{-6}}$ |
| $ch \times cond \times range$ | 19 | 0.84107 | 31.871 | $\mathbf{1.9207 \times 10^{-5}}$ |
